# NKX2.2 and KLF4 cooperate to regulate α cell identity

**DOI:** 10.1101/2024.08.07.607083

**Authors:** Elliott P. Brooks, McKenna R. Casey, Kristen L. Wells, Tsung-Yun Liu, Madeline Van Orman, Lori Sussel

## Abstract

Transcription factors (TFs) are indispensable for maintaining cell identity through regulating cell-specific gene expression. Distinct cell identities derived from a common progenitor are frequently perpetuated by shared TFs; yet the mechanisms that enable these TFs to regulate cell-specific targets are poorly characterized. We report that the TF NKX2.2 is critical for the identity of pancreatic islet α cells by directly activating α cell genes and repressing alternate islet cell fate genes. When compared to the known role of NKX2.2 in islet β cells, we demonstrate that NKX2.2 regulates α cell genes, facilitated in part by α cell specific DNA binding at gene promoters. Furthermore, we have identified the reprogramming factor KLF4 as having enriched expression in α cells, where it co-occupies NKX2.2-bound α cell promoters, is necessary for NKX2.2 promoter occupancy in α cells and co-regulates many NKX2.2 α cell transcriptional targets. Misexpression of *Klf4* in β cells is sufficient to manipulate chromatin accessibility, increase binding of NKX2.2 at α cell specific promoter sites, and alter expression of NKX2.2-regulated cell-specific targets. This study identifies KLF4 is a novel α cell factor that cooperates with NKX2.2 to regulate α cell identity.

## Introduction

The regulation of cell-specific expression by DNA binding transcription factors (TFs) is an integral part of establishing and maintaining cellular identity in all tissues. For cells with distinct functions that have differentiated from a common progenitor, preserving the unique gene expression that determines cellular identity is paramount. The establishment and maintenance of cell identity is often dependent on the function of cell-specific TFs that become activated during lineage specification and remain expressed throughout the lifetime of the cell. However, this simple paradigm can be challenged when a single TF regulates gene expression in multiple cell types that arise from a common lineage. Several mechanisms have been shown to be responsible for influencing the function of more broadly expressed TFs in a cell-context dependent manner, including: (1) RNA splice isoforms and alternative transcriptional start sites can produce cell type-enriched TF protein variants that possess distinct DNA binding affinities and result in cell-specific DNA occupancy profiles; (2) Post-translational modifications of TFs can occur in a cell type dependent manner causing altered physical interactions with DNA and/or co-factors; (3) Expression of cell-specific co-factors can interact with TFs to facilitate context dependent binding potentially by physically interacting with the TF and/or modulating local epigenetic characteristics of the chromatin. In an era where human stem cell therapies are becoming a reality, it is critical to understand the entire spectrum of mechanisms relating to the specification and maintenance of distinct cellular identities, including understanding how TFs expressed in a variety of cell types can acquire cell-specific functions.

The pancreatic endocrine cells represent an ideal model for studying the mechanisms of gene regulation by TFs because of the small number of functionally distinct cell-types that arise from a common endocrine progenitor. In the adult islet, there are four distinct hormone producing endocrine cell populations that arise from a common NEUROG3+ endocrine progenitor (Pictet et al., 1972, Smith et al., 2004, Prasadan et al., 2010, Bishop et al., 2013). In mice, α and β cells are the most numerous of the islet endocrine cell types, acting as a counterregulatory system to control blood glucose homeostasis. α cells secrete glucagon in response to hypoglycemia to stimulate the increase of blood glucose and β cells secrete insulin in response to hyperglycemia to stimulate the decrease of blood glucose. Until recently, research has predominantly focused on β cells due to their relatively straightforward connection with diabetes etiology and pathogenesis. However, there is growing appreciation that there are alterations in α cells in the context of type 1 and type 2 diabetes (T1D and T2D), in both mouse models and humans, including reduced α cell numbers, impaired glucagon secretion, and loss of identity that is characterized by misexpression of insulin and the dysregulation of TFs (Maclean and Ogilvie, 1959, Müller et al., 1970, Bolli et al., 1983, Gerich et al., 1973, Brissova et al., 2018, Marchetti et al., 2000, Dai et al., 2022, Brooks and Sussel, 2023). Since failure to maintain α cell identity is a potential etiological factor of diabetes, studying the molecular mechanisms of α cell identity maintenance is crucial to understanding the role of the whole islet in diabetes pathology.

Many studies in mice have identified cell-specific TFs that are essential for establishing and maintaining α cell identity and function, including Aristaless-related homeobox protein (ARX) and Musculoaponeurotic fibrosarcoma oncogene homolog B (MAFB) (Collombat et al., 2007, Courtney et al., 2013, Mastracci et al., 2011, Artner, 2006, Nishimura et al., 2006). However, the TFs Homeobox protein NK-2 homolog B (NKX2.2) and Paired box protein 6 (PAX6) are essential not only for α cell identity and function, but also essential for multiple endocrine lineages, including β cells (Sussel, 1998, Doyle and Sussel, 2007, Churchill et al., 2017, Gosmain et al., 2007, Gosmain et al., 2010, Swisa et al., 2017, Sander et al., 1997). Recent studies have demonstrated that PAX6 possesses unique transcriptional targets in α versus β cells due to the existence of distinct isoforms that influence cell-specific binding profiles (Singer et al., 2019). On the other hand, it is still not known how NKX2.2 exerts its cell-specific functions. NKX2.2 is expressed throughout pancreas development. In mice *Nkx2.2* is expressed in the PDX1+ pancreas progenitors, the NEUROG3 endocrine progenitor cell population and then becomes restricted to the α and β cell lineages, where expression is maintained throughout life (Arnes et al., 2012). Null mutations in *Nkx2.2* result in a complete lack of β cells and a severe reduction in α cells, demonstrating that it is essential for the formation of both islet cell lineages (Sussel, 1998). Furthermore, ablation of *Nkx2.2* from developing and adult β cells caused a decreased number of insulin expressing cells, impaired insulin secretion, and an increase in polyhormonal cells, including those that are converted to an α-like identity (Gutierrez et al., 2017). Transcriptomic analysis of *Nkx2.2*-ablated β cells supported the conclusion that in β cells, NKX2.2 directly activates β cell specific genes such as *Nkx6.1* and *Pdx1* while repressing α cell specific genes such as *Arx*. This suggests that β cells require NKX2.2 to maintain β cell identity and function by promoting the transcription of β cell enriched genes and repressing the expression of α cell enriched genes. The question then remains: If NKX2.2 is essential for repressing α lineage genes in β cells, what is the role of NKX2.2 in α cells?

To interrogate the role of NKX2.2 in α cells, we generated an α cell specific deletion of *Nkx2.2* in mice. Mice depleted for *Nkx2.2* in α cells displayed decreased fasted blood glucose levels, decreased α cell numbers, and remarkably, increased bihormonal cells. These results support a role for NKX2.2 in α cells that promotes α cell identity and represses alternate islet cell identities. Transcriptomic analysis of α cells with *Nkx2.2* ablation, suggested that NKX2.2 has opposing functions in α cells compared to β cells, by showing NKX2.2 activates many α cell specific genes while repressing many β and δ cell enriched genes. Genome occupancy studies to identify how NKX2.2 acquires these opposing functions revealed that NKX2.2 binds more frequently to proximal promoters in α cells versus its more frequent association with introns and putative enhancers in β cells. Finally, we identified Krueppel-like factor 4 (KLF4) as an α cell specific TF that influences the binding of NKX2.2 to promoter regions of the genome in α cells. Together, this study identifies a novel mechanism through which NKX2.2 adopts cell-specific functions in the pancreas and newly identifies KLF4 as an important factor in α cell development and function.

## Results

### Loss of Nkx2.2 in α cells causes a decrease in the α cell population and an increase in bihormonal cells

To understand the *in vivo* role of *Nkx2.2* in α cells we generated *Nkx2.2^fl/fl^;Gcg-iCre;Rosa26:tdTomato* (*Nkx2.2^αΔ^*) mice to constitutively knockout (KO) *Nkx2.2* within the α cell lineage (Shiota et al., 2017, Mastracci et al., 2013b). We observed high specificity of Cre activation in α cells from adult *Gcg-iCre;Rosa26:tdTomato* positive mice with respect to glucagon and insulin immunofluorescent (IF) labeling, consistent with previously published studies of the *Gcg-iCre* allele. Furthermore, the *Nkx2.2^αΔ^* KO mice showed a marked loss of NKX2.2 protein expression within the nuclei of tomato+ cells (Supplemental Fig. S1A). To determine the effect of *Nkx2.2* loss in α cells, we performed morphometric analysis of pancreatic tissue sections from 5-week-old (5wk) *Nkx2.2^αΔ^*mice. Both *Nkx2.2^+/+^;Gcg-iCre;Rosa26:tdTomato* (Gcg-iCre only) and *Nkx2.2^fl/fl^* (flox-only) mice were used as controls. Glucagon staining and visualization of the tdTomato reporter revealed a striking decrease in the size of the α cell population within the islet (Fig. 1A). Quantification of the glucagon expression area, as a measure of α cell area, revealed a ∼2.5-fold decrease in the mean glucagon expression area in the *Nkx2.2^αΔ^* compared to both the Gcg-iCre only and flox-only control mice (Fig. 1B). Quantification of glucagon+/tomato+ cells showed a 3-fold decrease in the average α cell number per islet in the *Nkx2.2^αΔ^* mice (Fig. 1C). These data suggest that NKX2.2 is important for producing and/or maintaining α cell population numbers.

**Figure 1.**
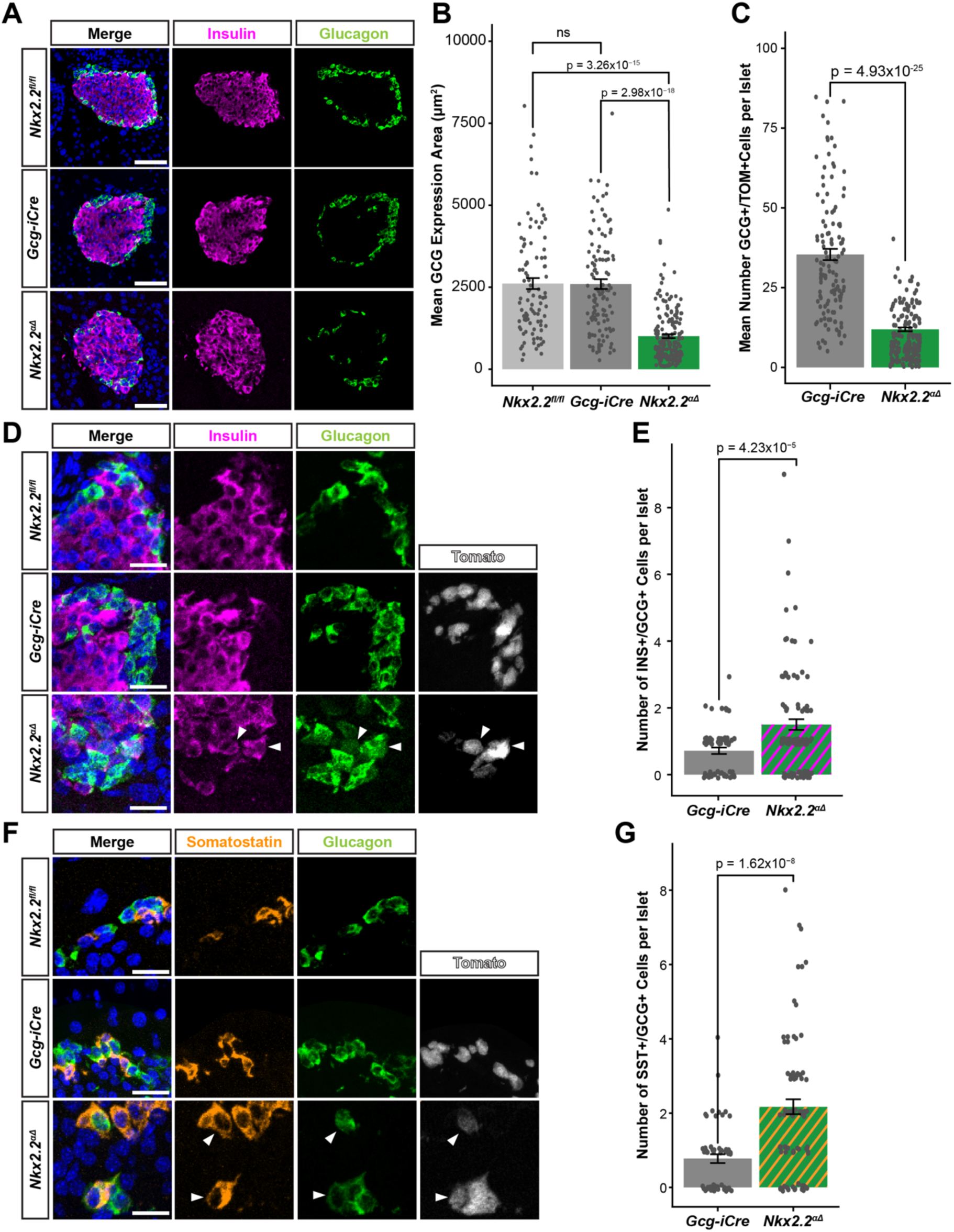
Knockout of *Nkx2.2* in α cell lineage causes decreased α cell population and hormonal dysregulation. (A) Glucagon (green), insulin (magenta) and DAPI (blue) expression in whole islets from 5wk *Nkx2.2^αΔ^* mutants and controls. Scale bar = 50 μm. (B) Quantification of mean glucagon expression area (μm^2^) for 5wk *Nkx2.2^αΔ^* mutants and controls. (C) Quantification of mean number of glucagon+/tomato+ cells per islet compared between 5wk *Nkx2.2^αΔ^* mutants and *Gcg-iCre* control. (D) Glucagon (green), insulin (magenta), tomato (white) and DAPI (blue) expression in 5wk *Nkx2.2^αΔ^* mutants and controls. White arrows point to insulin/glucagon bihormonal cells and scale bar = 20 μm. (E) Quantification of number of insulin/glucagon bihormonal cells per islet between *Nkx2.2^αΔ^* mutants and *Gcg-iCre* control. (F) Glucagon (green), somatostatin (orange), tomato (white) and DAPI (blue) expression in 5wk *Nkx2.2^αΔ^*mutants and controls. White arrows point to somatostatin/glucagon bihormonal cells and scale bar = 20 μm. (G) Quantification of number of somatostatin/glucagon bihormonal cells per islet for *Nkx2.2^αΔ^* mutants and *Gcg-iCre* control.

Previous studies showed that deletion of *Nkx2.2* specifically in the β cell lineage leads to the increase in insulin-β lineage cells and polyhormonal cells, including insulin/glucagon co-expressing cells (Papizan et al., 2011, Gutierrez et al., 2017). To assess whether *Nkx2.2* is also responsible for maintaining the distinct hormonal identity of α cells, we performed glucagon and insulin IF staining in pancreatic sections of 5wk *Nkx2.2^αΔ^* and control mice. This analysis demonstrated that the number of cells that co-expressed both glucagon and insulin were present in a higher frequency in *Nkx2.2^αΔ^* compared to Cre control islets (Fig. 1D,E). Similarly, there was a significant increase in the frequency of glucagon+/somatostatin+ bihormonal cells in the *Nkx2.2^αΔ^* mutant islets (Fig. 1F,G). Although an expansion of ghrelin expressing cells was seen in the global NKX2.2 KO mouse islets, we did not detect an increase in ghrelin+/glucagon+ bihormonal cells (Supplemental Fig. S1B). Together, this analysis indicates that NKX2.2 regulates the population size of α cells and represses insulin and somatostatin expression in the α cells that remain. Interestingly, this suggests that NKX2.2 is simultaneously able to maintain the opposite cell fates of two closely related islet lineages.

### NKX2.2 regulates the α cell response to fasting

A fundamental function of α cells is to secrete glucagon in response to low glycemic levels. To examine whether depletion of NKX2.2 influences α cell function, we measured the *ad libitum* blood glucose levels at 4 wk. Like many other animal models with α cell dysfunction or lack of α cells, there was no significant difference in blood glucose between the *Nkx2.2^αΔ^* mice and the Gcg-iCre control mice in basal conditions (Supplemental Fig. S1C)(Heller et al., 2004, Wilcox et al., 2013, Conrad et al., 2016). Furthermore, we observed no difference in body mass between the KO and control mice (Supplemental Fig. S1C). To assess the response of *Nkx2.2^αΔ^* mice under metabolic stress, mice were fasted for 6 hours. The *Nkx2.2^αΔ^* mice had a significantly lower mean fasted blood glucose level than the combined mean of the flox-only and Gcg-iCre only control mice (Supplemental Fig. S1D). However, in response to insulin induced hypoglycemia, the 5wk *Nkx2.2^αΔ^*fasted mice only trended towards blunted insulin tolerance (Supplemental Fig. S1D), suggesting that, again consistent with other mouse models harboring α cell defects, loss of NKX2.2 only has a subtle effect on α cell functions.

### NKX2.2 regulates cell-specific gene expression in α cells

Previous studies assessing the regulatory role of *Nkx2.2* in β cells demonstrated that NKX2.2 is essential for activating β cell lineage genes and repressing non-β cell islet lineage genes (Churchill et al., 2017, Doyle et al., 2007, Doyle and Sussel, 2007, Gutierrez et al., 2017, Papizan et al., 2011, Sussel, 1998). To determine the NKX2.2 transcriptional targets in α cells, we performed global transcriptome analysis on Fluorescent Activated Cell Sorted (FACS) α cells from *Nkx2.2^αΔ^* mouse islets versus Gcg-iCre only control islets based on tomato reporter expression. Enrichment of the α cell population in the tomato+ fraction was validated by qPCR for expression of *Gcg* and *Ins2* mRNA (Supplemental Fig. S2A,B,C). Consistent with the morphometric analysis, there was a statistically significant decrease in the percentage of α cells collected from the *Nkx2.2^αΔ^* mice compared to the Gcg-iCre-only control (Supplemental Fig. S2D). RNA-sequencing and differential gene expression (DGE) analysis revealed that deletion of *Nkx2.2* in α cells resulted in 1,371 differentially expressed genes (DEGs) of which 669 were downregulated and 702 were upregulated (Supplemental Data 1). To identify the genes dysregulated in the *Nkx2.2^αΔ^* mice that were known to have enriched expression in either α, β or δ cell populations, we then performed a pair-wise DGE analysis utilizing a previously published RNA-seq data set from sorted wildtype mouse α, β and δ cells (DiGruccio et al., 2016). This analysis revealed that loss of *Nkx2.2* in α cells resulted in the downregulation (de-activation) of 160 α cell specific genes, including *Gcg, MafB, Fev, Slc38a5, Irx2* and *Etv1*, and upregulation (de-repression) of 66 β cell specific genes, including *MafA, Pcsk1 (Pc1), Ucn3, Slc2a2 (Glut2), Ins1* and *Ins2* (Fig. 2A). We also identified 108 δ cell genes that became upregulated (de-repressed) in the *Nkx2.2^αΔ^* α cells. To compare how NKX2.2 regulates cell-specific gene expression in α versus β cells we conducted hypergeometric enrichment analyses of cell-specific gene expression in the *Nkx2.2^αΔ^*RNA-seq and in a previously published *Nkx2.2* β-KO RNA-seq (Fig. 2B)(Gutierrez et al., 2017). These analyses revealed that in the *Nkx2.2^αΔ^*, α cell specific genes were enriched in the downregulated gene set, whereas β cell specific genes were enriched in the upregulated gene set. This is in direct contrast to the *Nkx2.2* β-KO mice in which α genes were enriched in the upregulated gene set and β genes were enriched in the downregulated gene set. As expected, δ cell genes were enriched in the gene sets that were upregulated by both the *Nkx2.2^αΔ^* and *Nkx2.2* β-KO. These analyses suggest that NKX2.2 functions in α cells to activate α cell genes and represses β cell genes, whereas it has opposite gene regulatory functions in β cells.

**Figure 2.**
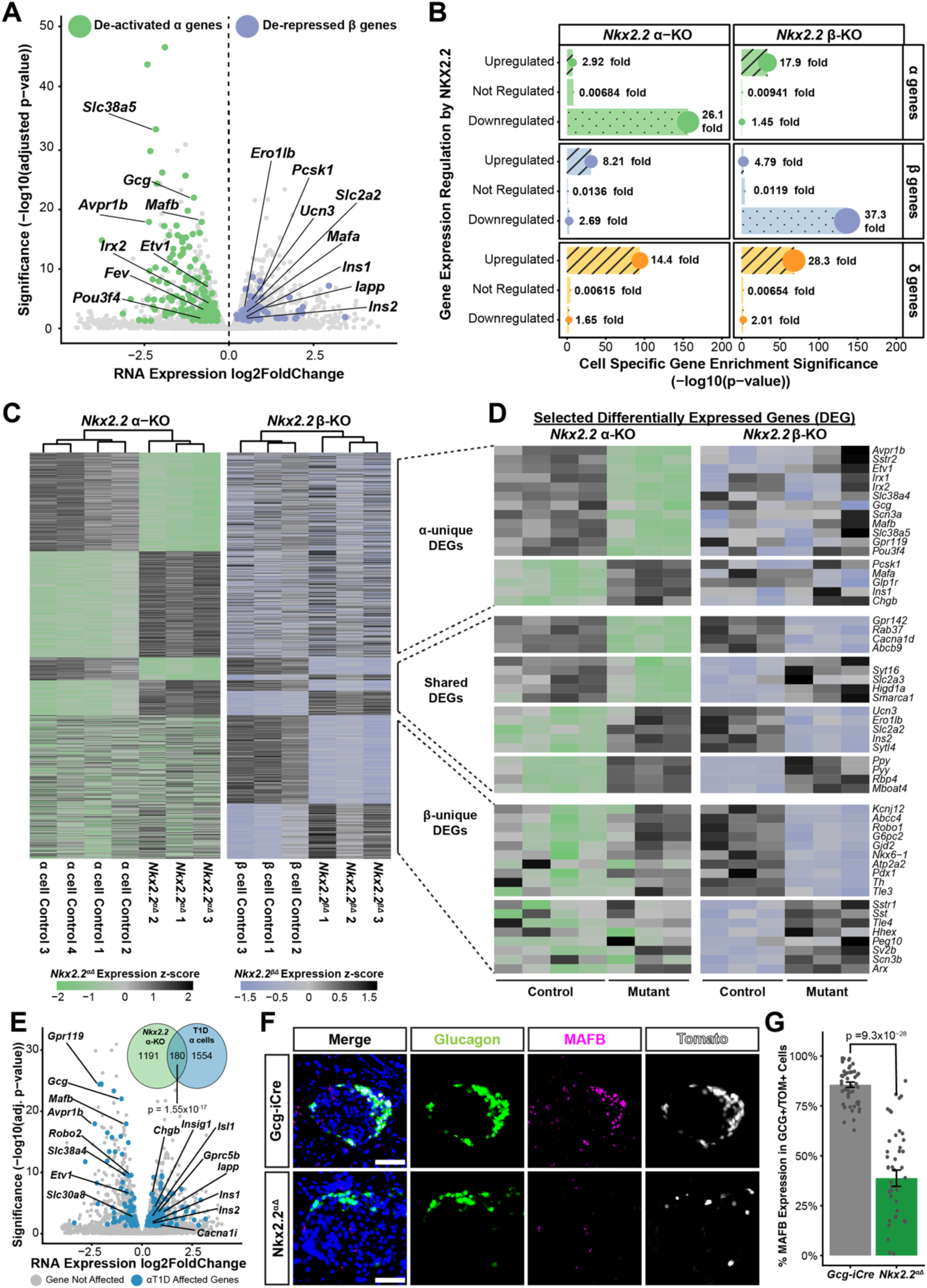
NKX2.2 has context-dependent transcriptional targets in α and β cells. (A) Volcano plot showing the log2 foldchange and significance of DGE found in *Nkx2.2^αΔ^*α cell RNA-seq—highlighting the presence of de-activated α genes (green) and de-repressed β genes (purple). (B) Hypergeometric gene-set enrichment testing showing statistical significance for enrichment of α (green), β (purple) and delta (orange) expressed genes within upregulated genes (hashed-bar) and downregulated genes (dotted-bar) from the *Nkx2.2*-αΚΟ and published *Nkx2.2*-βΚΟ RNA-seqs. Size of point denotes foldchange over background. (C) Heatmap of expression z-scores for all DEGs found in both the *Nkx2.2*-αΚΟ and published *Nkx2.2*-βΚΟ RNA-seqs. Rows aligned by gene for all samples and columns organized by non-biased clustering. (D) Expression z-score heatmap of selected DEGs representative of NKX2.2 α-unique targets, NKX2.2 β-unique targets and NKX2.2 targets shared in α and β cells. (E) Volcano plot of *Nkx2.2^αΔ^* α cell RNA-seq DEGs overlapped with published genes dysregulated in α cells with T1D. Inset Venn diagram represents the overlap statistics between RNA-seq datasets. (F) Glucagon (green), MAFB (magenta), tomato (white) and DAPI (blue) expression in 5wk *Nkx2.2^αΔ^*mutants and *Gcg-iCre* control. Scale bar = 50 μm. (G) Quantification of percentage of glucagon+/tomato+ α cells that express MAFB protein in *Nkx2.2^αΔ^* mutants and *Gcg-iCre* control islets.

To identify the cell type-enriched genes that composed NKX2.2 cell-specific transcriptional targets, we compared the α and β cell *Nkx2.2* KO DEGs (Fig. 2C). Among the 1,116 DEGs present only in the *Nkx2.2^αΔ^* mice were many genes encoding TFs (*Etv1, Irx1/2, MafB, Pou3f4*), ion channels related to endocrine function (*Kcnk3, Kcnc4, Cacng4, Scn3a*), α cell enriched cell surface receptors (*Avpr1b, Gpr119, Sstr2*), and solute channels related to α cell proliferation (*Slc38a5, Slc38a4, Slc7a2)(*Fig. 2D). Among the 706 DEGs found only in the β cell *Nkx2.2* KO mice were those encoding essential β cell TFs (*Nkx6.1, Pdx1*), genes related to intracellular signaling (*Atp2a2, Atp2a3, Pde5a*), insulin and secretory vesicle related genes (*G6pc2*), and ion channels (*Abcc4, Kcnj12, Scn1b*). Several genes were regulated by NKX2.2 in both cell types, including a subset of genes that are oppositely regulated in α and β cells, supporting the ability of NKX2.2 to switch regulatory roles on the same gene in a cell-context dependent manner. Together these comparative analyses support the notion that *Nkx2.2* has α and β cell specific molecular functions that depend on cellular context.

### NKX2.2 in α cells regulates genes affected in T1D α cells

Recent studies have demonstrated there is transcriptional dysregulation in α cells isolated from T1D individuals, including the downregulation of *Nkx2.2* (Brissova et al., 2018). To determine the proportion of genes dysregulated in T1D α cells that were potentially transcriptional targets of NKX2.2 in α cells, we reanalyzed the published α cell T1D RNA-seq via DGE analysis and intersected those DEGs with the DEGs found in our *Nkx2.2^αΔ^*mouse RNA-seq. Interestingly, there was a statistically significant overlap of 180 homologous genes in α cells that were dysregulated by both T1D in humans and *Nkx2.2^αΔ^* in mice (Fig. 2E). This included the downregulation of Gcg, *MafB, Avpr1b* and *Etv1* and the upregulation of *Ins1, Ins2,* and *Isl1*. Since *MafB* is known for its critical role in α cell maturation and maintenance, we validated the downregulation of *MafB* expression in the *Nkx2.2^αΔ^*mice. IF analysis on 5wk mouse pancreas revealed that *Nkx2.2^αΔ^*α cells have a ∼50% decrease in tomato+ α cells expressing MAFB (Fig. 2F,G).

### NKX2.2 has cell-specific DNA binding and binds more to proximal promoters in α cells

NKX2.2 is a homeodomain TF that regulates islet cell transcription via direct DNA binding (Doyle et al., 2007, Gutierrez et al., 2017, Mastracci et al., 2013a, Papizan et al., 2011). This motivated the question of whether the distinct α and β cell specific transcriptional targets of NKX2.2 are due to cell-specific DNA occupancy. Genome wide DNA binding occupancy of NKX2.2 in α cells identified by chromatin immunoprecipitation coupled with sequencing (ChIP-seq) in the αTC α cell line uncovered 7,389 NKX2.2 genomic binding regions (Fig. 3A). Integration of the “nearest genes and genomic feature” annotations of the NKX2.2 α cell ChIP peaks with the DEGs found in the *Nkx2.2^αΔ^* RNA-seq revealed that NKX2.2 binds to 1,101 loci associated with genes that are dysregulated in *Nkx2.2^αΔ^* α cells (Fig. 3A), including important islet cell genes such as *MafB, Irx2, Etv1, Fev, Pcsk1*, *Ero1lb*, and *Iapp (Yu et al., 2015)*.

**Figure 3.**
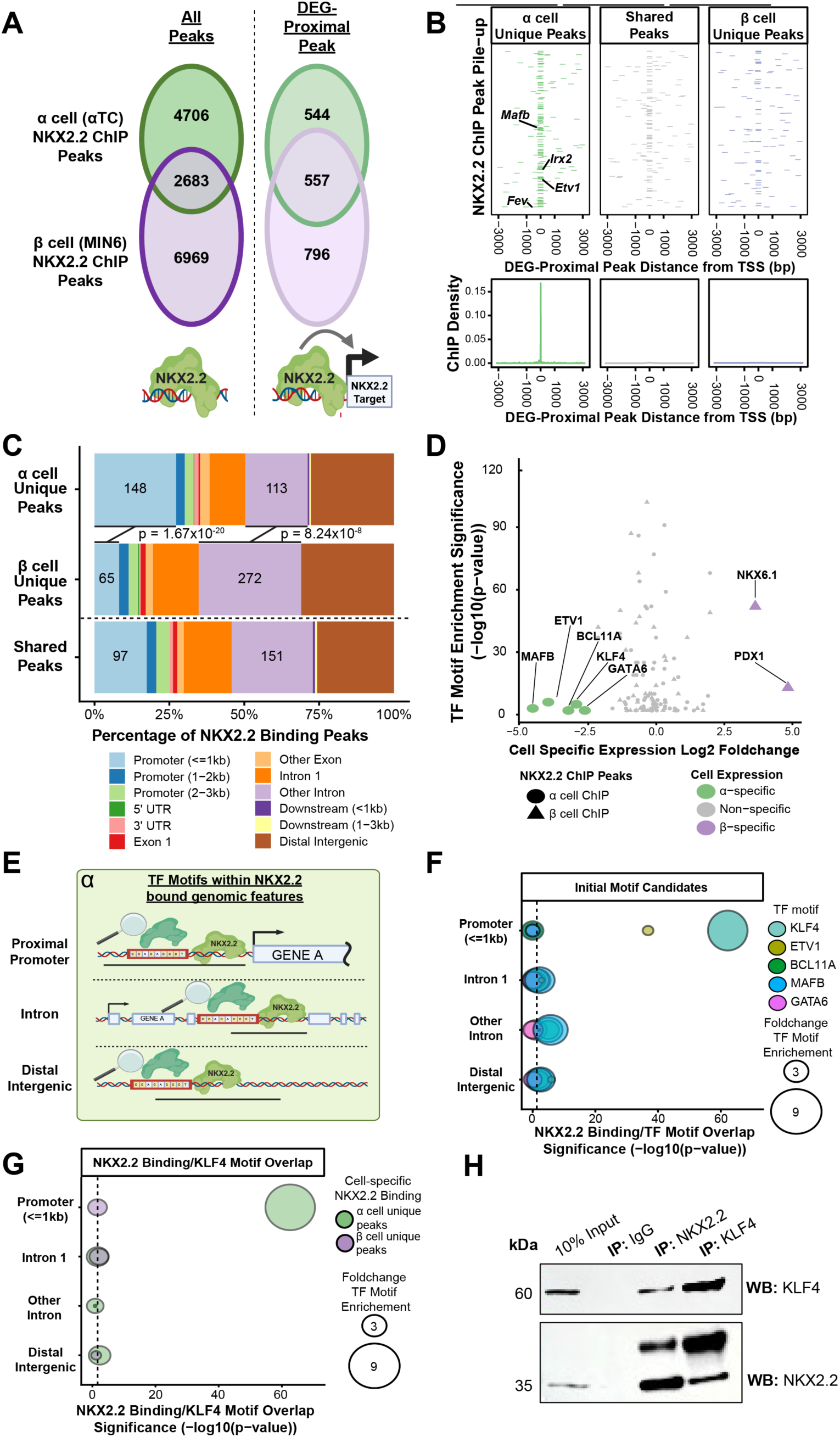
NKX2.2 has α cell specific genome binding that is enriched for KLF4 binding motifs. (A) Venn diagrams showing the genomic overlap of NKX2.2 ChIP binding sites in α cells and β cells for all binding sites and sites that are associated with gene expression changes due to *Nkx2.2*-KO. (B) Upper plots: Pile-up plots for the distance of promoter-associated direct target NKX2.2 ChIP peaks to the transcriptional start sight; subset by α/β-unique binding sites and shared binding sites. Lower plots: density of NKX2.2 ChIP pile-up near transcriptional start sites. (C) Distribution of direct target NKX2.2 binding sites at key genomic annotations; subset by α/β-unique binding sites and shared binding sites. Numbers in bars represent quantity of peaks in those genomic annotations. (D) Volcano plot showing the α-vs-β cell DEGs and enrichment significance of TFs binding motifs around NKX2.2 binding sites. (E) Schematic illustrating our motif analysis paradigm that searches for motifs for α cell enriched TFs near promoter, intron and distal intergenic loci bound by NKX2.2 in α cells. (F,G) Dot plots showing the significance of motif enrichment of (F) initial α cell enriched TF candidates within α cell NKX2.2 binding sites and (G) KLF4 motif enrichment near α-unique and β-unique binding sites. Size of dot represents the foldchange enrichment over genomic background frequency and dotted line represents log transformed p-value cutoff of 0.05 (H) Representative Western blot of NKX2.2 and KLF4 co-IPs in αTC cell culture. Input and IgG-IP included as positive and negative controls, respectively.

To ascertain if NKX2.2 binding differs between α and β cells, we compared NKX2.2 α cell ChIP-seq to a previously published β cell NKX2.2 ChIP-seq performed in MIN6 β cells (Gutierrez et al., 2017). Here, we defined overlap of α and β NKX2.2 binding peaks as having at least 1 bp overlap within +/- 100 bp of the ChIP peak summit. Remarkably, this analysis identified only 19% of all NKX2.2 binding sites are shared between α and β cells (Fig. 3A). Furthermore, NKX2.2 occupied 4,706 sites unique to α cells and 6,969 sites unique to β cells. In addition, overlap analysis of direct target NKX2.2 binding peaks in α and β cells found that 71% of all direct target binding sites of NKX2.2 are unique to either α or β cells (Fig. 3A). This suggests that NKX2.2 regulates transcription in a cell-specific manner in part by binding to genomic regions that are unique to α or β cells.

We further characterized the DNA binding of NKX2.2 in α versus β cells by plotting the pile-up of NKX2.2 binding peaks in relation to the nearest transcriptional start site (TSS). Strikingly, NKX2.2 genomic occupancy that is unique to α cells had the highest percentage of TSS-proximal peak pile-up and density compared to shared or β cell unique peaks, which is especially apparent when only examining direct target associated binding (Supplemental Fig. S3A; Fig 3B). The disproportionate occupancy of NKX2.2 at promoter regions in α cells motivated the investigation of the general genome wide distribution of NKX2.2 binding sites in α and β cells. This analysis identified a highly significant increase in α cell unique NKX2.2 occupancy at promoter regions located ≤ 1kb from the TSS when compared to occupancy sites that were unique to β cells (Fig. 3C; Supplemental Fig. S3B). Alternatively, regions of NKX2.2 occupancy that were unique to β cells were predominantly found in “other intron” (excludes 1^st^ introns) regions. Of note, NKX2.2 was bound to the proximal promoters of the α cell enriched genes *MafB, Irx2, Fev* and *Etv1* in α cells but not in β cells (Fig. 3B). Together these findings suggest that a large proportion of the difference in NKX2.2 transcriptional targets between α and β cells is characterized by increased NKX2.2 occupancy at proximal promoters in α cells versus putative enhancer regions in β cells.

### KLF4 motifs are enriched in NKX2.2-bound proximal promoters in α cells

The data suggests that NKX2.2 regulates transcription in a cell-context dependent manner by binding to cell-specific genomic regulatory regions. Currently, there is no evidence to suggest that NKX2.2 has any cell-specific isoforms or post-translational modifications that could account for a biochemical change in the binding specificity of NKX2.2 between α and β cells. TFs often regulate transcription of target genes through cooperative DNA binding. Indeed, post-hoc multi-omic analysis of TF binding and chromatin accessibility has identified many islet putative enhancers and functional loci that are occupied by multiple TFs, including NKX2.2 (Mawla et al., 2023). Thus, we hypothesized that a TF with α cell enriched expression would bind closely to NKX2.2 at α cell proximal promoters to facilitate α cell specific NKX2.2 binding. To identify potential TF co-occupancy candidates, we performed a non-biased search for enriched TF binding motifs located proximally to α or β NKX2.2 occupancy sites using the HOMER2 software and motif database (Heinz et al., 2010). We then intersected islet specific gene expression with the enriched motifs to determine an association with α or β cell specific gene expression (DiGruccio et al., 2016). This analysis identified 5 motifs that corresponded to α cell enriched TFs (MAFB, ETV1, BCL11A, KLF4, and GATA6) and were only enriched at regions occupied by NKX2.2 in α cells. It also identified 2 motifs that corresponded to β cell enriched TFs (NKX6.1 and PDX1) that were preferentially enriched at NKX2.2 binding sites in β cells (Fig. 3D). To further validate the 5 α-enriched TFs as candidates for cooperation with NKX2.2 we characterized the enrichment of their respective motifs specifically at the genomic features (proximal promoters, introns, and distal intergenic regions) differentially occupied by NKX2.2 in α versus β cells (Fig. 3E). This analysis identified KLF4 and ETV1 motifs as significantly enriched in proximity to NKX2.2 binding sites at proximal promoters in α cells, with KLF4 having the greatest and most significant promoter enrichment (Fig. 3F). We further assessed the specificity of KLF4 motif enrichment in α cell promoters bound by NKX2.2 by performing KLF4 motif analysis on NKX2.2 binding sites that were unique to either α or β cells. Interestingly, KLF4 motifs were only enriched in proximal promoters with α-unique NKX2.2 binding, but not in promoters with β-unique NKX2.2 binding or at other genomic locations (Fig. 3G, Supplemental Data 2). We further characterized the biochemical relationship between NKX2.2 and KLF4 in α cells by performing co-immunoprecipitation (co-IP) of endogenous NKX2.2 or endogenous KLF4 in αTC cells. An interaction between NKX2.2 and KLF4 protein was detected both in anti-KLF4 co-IPs and anti-NKX2.2 co-IPs, suggesting that NKX2.2 and KLF4 physically interact in α cells (Fig. 3H). Together these data suggest that KLF4 could be co-occupying the proximal promoters that are uniquely bound by NKX2.2 in α cells.

### MafB and Irx2 promoters are co-occupied by NKX2.2 and KLF4 in α cells

Both MAFB and IRX2 are prominent α cell enriched TFs in mice and humans and several studies have shown that the TF MAFB is important for the maturation and maintenance of α cells (Artner, 2006, Chang et al., 2020, Chung et al., 2010, Conrad et al., 2016, Katoh et al., 2018). In this study we demonstrate that NKX2.2 in α cells activates the expression of *MafB* and *Irx2*, motivating us to further investigate the NKX2.2 regulatory mechanism of these genes. NKX2.2 ChIP-seq performed in the αTC cells vs. MIN6 cells revealed that NKX2.2 occupies the proximal promoters of *MafB* and *Irx2* only in α cells (Fig. 4A,B). Furthermore, the assay for transposase accessible chromatin using sequencing (ATAC-seq) in αTCs and MIN6 cells showed that chromatin is open at the *MafB* and *Irx2* promoters in α cells but not β cells, supporting the α cell specific activity of these promoters. We performed Cleavage Under Targets and Release Using Nuclease (Cut&Run) qPCR to validate that the promoters of *MafB* and *Irx2* were bound by NKX2.2 in α cells. Using primers designed to amplify the region surrounding the NKX2.2 ChIP-seq peak at the *MafB* or *Irx2* promoters we found that NKX2.2 Cut&Run samples were highly enriched over the IgG negative control, confirming NKX2.2 occupancy at these loci (Fig. 4C,B). Furthermore, motif analysis identified several highly complementary KLF4 motifs located near the NKX2.2 binding site at the *MafB* and *Irx2* promoters, suggesting KLF4 co-occupancy. To test the potential binding of KLF4 at these loci we performed Cut&Run qPCR using primers that amplify the NKX2.2-bound region of the *MafB* and *Irx2* promoters. KLF4 immuno-enriched Cut&Run samples were found to have DNA enrichment at the *MafB* and *Irx2* promoters compared to the IgG control. Together, these findings suggests that NKX2.2 and KLF4 co-occupy the proximal promoters of *MafB* and *Irx2* in α cells.

**Figure 4.**
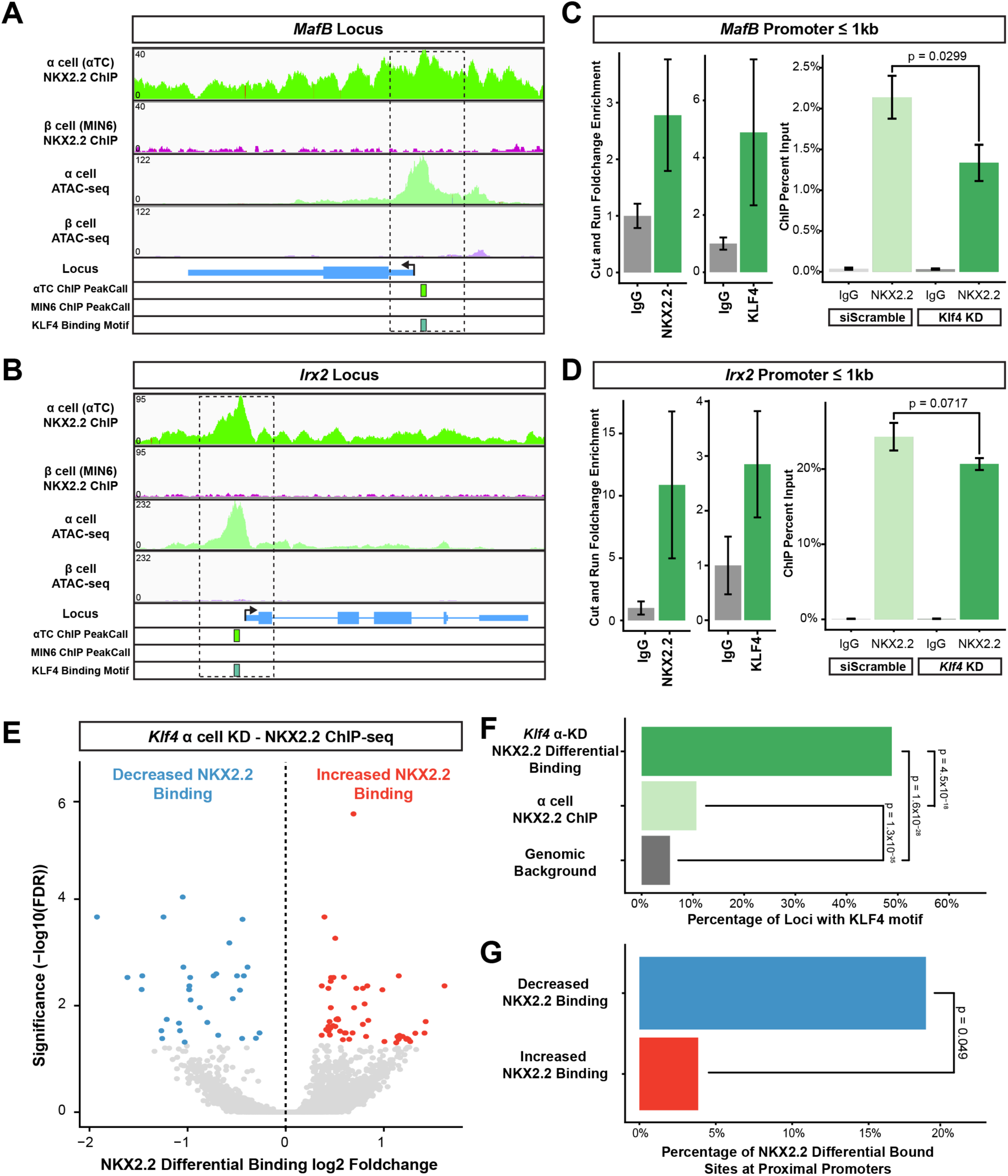
KLF4 co-occupies α cell specific NKX2.2 binding sites and depletion of KLF4 causes NKX2.2 binding dysregulation in α cells. (A,B) Genomic views at the *MafB* and *Irx2* loci showing the NKX2.2 α and β cell ChIP-seq, α and β cell ATAC-seq, ChIP peak-calling and KLF4 motif presence. Dotted boxes outline the *MafB* or *Irx2* transcriptional start sites. (C,D) Left: NKX2.2 and KLF4 binding at the *MafB* and *Irx2* proximal promoters of α cells. Right: NKX2.2 binding at the *MafB* and *Irx2* proximal promoters for siScramble control and *Klf4*-KD α cells. (E) Volcano plots showing the differential binding of NKX2.2 caused by *Klf4*-KD in α cells. (F) Plot showing the percentage of *Klf4*-KD differential NKX2.2 binding sites harboring a KLF4 binding motif and compared to the KLF4 motif presence at background levels and all NKX2.2 α cell binding sites. (G) Percentage of NKX2.2 differential binding sites found at proximal promoters and subset by decreased and increased NKX2.2 binding.

### KLF4 is required to maintain the binding of NKX2.2 at α cell specific binding sites

The data suggests that KLF4 occupies loci that are bound by NKX2.2 in α cells, but not β cells. Due to its known function as a pioneer TF, we hypothesized that KLF4 is facilitating the cell-specific binding and subsequent cell-specific transcriptional regulation performed by NKX2.2 in α cells (Nakagawa et al., 2008, Takahashi and Yamanaka, 2006, Soufi et al., 2012). To test this hypothesis, we assessed whether NKX2.2 occupancy at the *MafB* and *Irx2* proximal promoters were affected by knocking down (KD) *Klf4* expression in αTC cells (Supplemental Fig. S4A). As expected from the global ChIP-Seq analysis, NKX2.2 binding at the *MafB* and *Irx2* promoters was highly enriched at both promoters in the siScramble control αTC cells. However, NKX2.2 binding at the *MafB* promoter was significantly reduced by 38% in the *Klf4*-KD αTC cells (Fig. 4C). There was also a trending 15% decrease in NKX2.2 binding at the *Irx2* promoter, although it did not reach significance (Fig. 4D). These results suggest that KLF4 is required to maintain proper occupancy levels of NKX2.2 at the *MafB* promoter. To test if KLF4 can broadly facilitate NKX2.2 binding, we performed an NKX2.2 ChIP-seq in *Klf4*-KD α cells. A differential binding analysis found that depletion of *Klf4* resulted in the differential binding of NKX2.2 at 84 loci, including both a decrease and increase of NKX2.2 binding (Fig. 4E). We searched for KLF4 binding motifs within the NKX2.2 differentially bound sites caused by the *Klf4*-KD and found that 48% of all NKX2.2 differential binding sites contained a high-scoring KLF4 motif, a statistically significant enrichment of KLF4 motifs over genomic background. Furthermore, NKX2.2 differentially bound sites were significantly more enriched for KLF4 motifs than all NKX2.2 α cell binding sites (Fig. 4F). Genomic annotations of the NKX2.2 differentially bound sites also revealed a higher percentage of NKX2.2 binding loss at proximal promoters (Fig. 4G, Supplemental Fig. S4B). This suggests that the NKX2.2 differentially bound sites that were observed following depletion of *Klf4* in α cells is likely due to the proximal binding of KLF4 and suggests that KLF4 facilitates the binding of NKX2.2 in α cells, especially by increasing binding at promoters.

### KLF4 and NKX2.2 co-regulate gene expression in α cells

KLF4 is a well characterized reprogramming factor that is able to manipulate chromatin accessibility, histone modification, and 3D chromatin structure as a means to regulate transcriptional targets (Di Giammartino et al., 2019, Moonen et al., 2022, Rosenzweig et al., 2013, Wei et al., 2013). Importantly, *Klf4* expression within the islet is enriched in α cells versus β cells, but the role of KLF4 in α cells has not been explored previously (DiGruccio et al., 2016, Schaum et al., 2018). To determine whether NKX2.2 and KLF4 possess overlapping transcriptional targets in α cells, we used siRNA to knockdown KLF4 in αTC cells and performed RNA-seq analysis (Supplemental Fig. S5A, Supplemental Data 3). The *Klf4*-KD cells had dysregulation of 1,330 genes compared to the siScramble control (Supplemental Fig. S5B) of which 209 genes (16%) were also differentially expressed in *Nkx2.2^αΔ^*α cells, including the mutual downregulation of α-enriched genes *MafB* and *Slc41a2* and upregulation of the β gene *Ero1lb*, further suggesting that KLF4 and NKX2.2 cooperate to regulate α cell identity.

To interrogate the role of KLF4 in α cells *in vivo* we then generated *Klf4^fl/fl^;Gcg-iCre* (*Klf4^αΔ^*) mice to constitutively delete *Klf4* in the α cell lineage (Yoshida et al., 2008). Morphometric analysis of the mutant islets revealed that there was no significant difference in α cell numbers between *Klf4^αΔ^* and *Gcg-iCre* control 5wk islets (Fig. 5A, B, Supplemental Data 4). However, transcriptomic analysis of *Klf4^αΔ^* islets vs. *Gcg-iCre* control islets isolated from 5wk mice revealed there was significant dysregulation of nearly 400 genes, including the downregulation of α-enriched *genes Gcg, Arx, Slc38a5* and *Irx2* and a trending downregulation of α-enriched genes *MafB, Fev*, and *Etv1* (Fig. 5C). In addition, there was an upregulation of several β cell genes, including *Sytl4, Gcgr* and *Tle3*. Intersection of the *Klf4^α^*^Δ^ islet DEGs with *Nkx2.2^αΔ^* DEGs found that 13% of the dysregulated genes in *Klf4^αΔ^* were also dysregulated in *Nkx2.2^αΔ^*. Importantly, 48 high-confidence NKX2.2/KLF4 overlapping transcriptional targets included the α-enriched genes *Gcg, Irx2, Higd1a, Gpr119* and *Slc38a5* and β-enriched genes *Sytl4* and *Diras2* (Fig. 5D). Further integration of the genes that were differentially expressed *in vitro* and *in vivo* determined that 25% of the 48 high-confidence NKX2.2/KLF4 *in vivo* DEGs were also dysregulated by *Klf4*-KD *in vitro* (Supplemental Fig. S5B). Together these analyses support the co-regulation of the α cell transcriptome by KLF4 and NKX2.2 and suggest for the first time that KLF4 is required for maintenance of the α cell lineage transcriptional identity.

**Figure 5.**
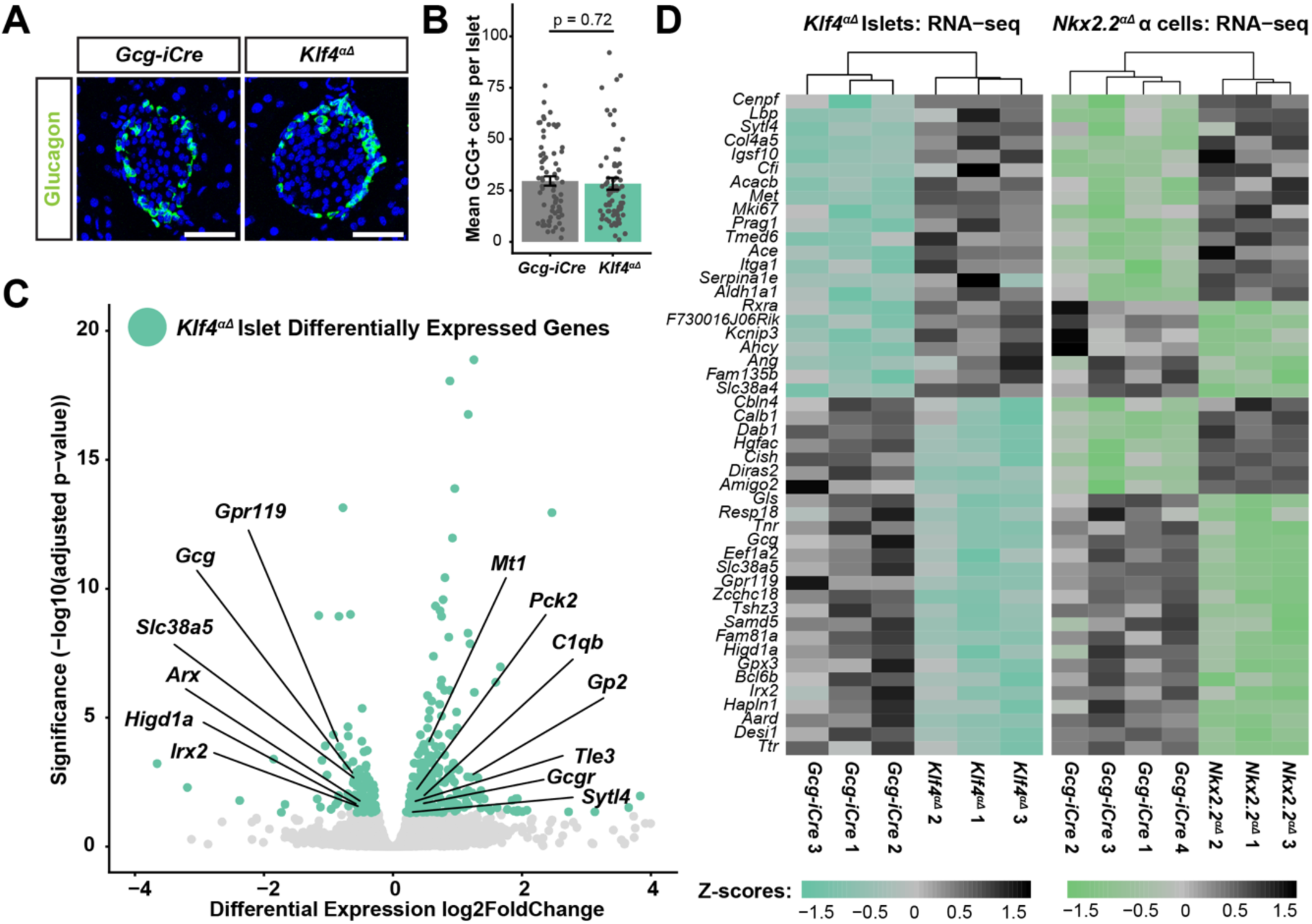
KLF4 and NKX2.2 co-regulate the α cell transcriptome. (A) Glucagon (green) and DAPI (blue) expression for 5wk *Klf4^αΔ^* mutants and *Gcg-iCre* controls. (B) Quantification of the number of glucagon+ cells per islet for 5wk *Klf4^αΔ^*mutant and *Gcg-iCre* control islets. (C) Volcano plot showing the significance and log2 foldchange of genes dysregulated by *Klf4*-KO in α cells. (D) Expression z-score heatmap of genes found to be dysregulated in both the *Klf4^αΔ^* mutants and *Nkx2.2^αΔ^* mutants. Rows aligned by gene for all samples and columns organized by non-biased clustering.

### Klf4 misexpression in β cells manipulates chromatin accessibility at NKX2.2 cell-specific binding sites

To test the sufficiency of the KLF4 interaction to potentiate the α cell unique genomic occupancy of NKX2.2, we misexpressed myc-tagged *Klf4* in MIN6 cells (*Klf4*-OE)(Supplemental Fig. S6A)(Mourao et al., 2021). Differential chromatin accessibility analysis using ATAC-seq revealed that *Klf4*-OE in MIN6 β cells caused 55,838 chromatin regions to increase or decrease significantly in accessibility (Fig. 6A, Supplemental Data 5). Interestingly, integrative analysis of the differential chromatin accessibility with cell-specific NKX2.2 binding sites found that *Klf4*-OE in β cells caused differential chromatin accessibility at 51% of all α-unique NKX2.2 binding loci and 43% of all β-unique NKX2.2 binding loci (Fig. 6A). Based on our hypothesis that KLF4 mediates the cell-specific binding of NKX2.2 in α cells, we postulated that misexpression of *Klf4* in β cells would have 2 major outcomes: (1) “pro-α” conditions where *Klf4*-OE caused an increase in chromatin accessibility that overlapped with an α-unique NKX2.2 binding site and (2) “anti-β” conditions where *Klf4*-OE caused a decrease in chromatin accessibility that overlapped with a β-unique NKX2.2 binding site (Fig. 6B). In other words, KLF4 promotes the NKX2.2 α cell binding profile in α cells by opening chromatin at α cell promoters but antagonizes NKX2.2 binding at β-unique loci by closing chromatin. Consistent with this hypothesis, 58% of cell-specific NKX2.2 binding sites that overlap with a *Klf4*-OE differential chromatin site corresponded to the pro-α or anti-β paradigm, including the proximal promoters of α-enriched genes *Fev, Irx1, Irx2* and *MafB* and the regulatory regions of β cell genes, *Mafa, Ins2, Nkx6.1* and *Ins1* (Fig. 6C). These data together show that *Klf4* misexpression in β cells is sufficient to open chromatin at many α cell specific NKX2.2 binding sites and close chromatin at many β cell specific NKX2.2 binding sites. This supports the model that KLF4 promotes a chromatin landscape that is conducive to the α cell specific NKX2.2 binding profile and antagonistic to the β cell binding profile of NKX2.2.

**Figure 6.**
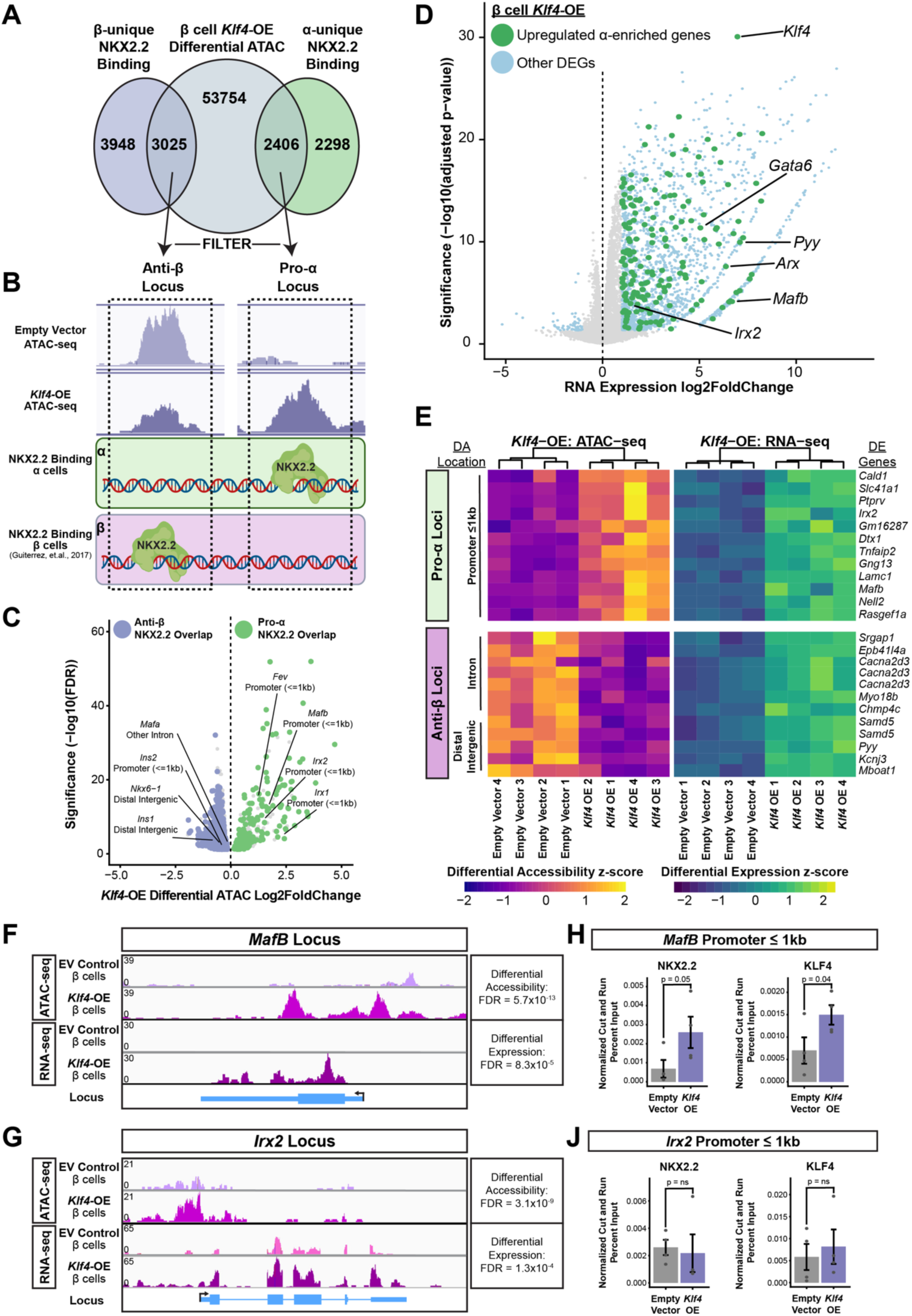
Misexpression of *Klf4* in β cells causes changes to chromatin accessibility, gene expression and NKX2.2 binding that resemble α cell profiles. (A) Overlap of cell-specific NKX2.2 binding sites with differential chromatin accessibility sites caused by *Klf4*-OE in β cells. (B) Schematic illustrating the “pro-α/anti-β” hypothesis-driven filtering paradigm of β cell *Klf4*-OE differential ATAC-seq sites that overlap with cell-specific NKX2.2 binding sites. (C) Volcano plot of the significance and log2 foldchange for all differential ATAC sites that overlap with NKX2.2 binding sites with “pro-α/anti-β” overlap conditions denoted by color. (D) Volcano plot of significance and log2 foldchange of DEGs found in RNA-seq of *Klf4*-OE β cells. Genes known to be α cell-enriched are labeled in green. (E) Heatmaps of selected NKX2.2 binding site loci that overlap with differential chromatin accessibility and associated gene expression change caused by *Klf4*-OE in β cells. Whether the loci are a “pro-α” or “anti-β” condition and the genomic feature annotation are labeled on the left and associated gene names are labeled on the right. (F, G) Sequencing read pile-up plots at the *MafB* and *Irx2* Loci for the ATAC-seq and RNA-seq experiments done in MIN6 cells with *Klf4-*OE or empty vector conditions. (H,J) Binding assays measuring the enrichment of NKX2.2 and KLF4-myc bound DNA at the *MafB* and *Irx2* promoters for the empty vector controls and *Klf4*-OE MIN6 cells.

### β cell misexpression of Klf4 causes transcriptional changes associated with differential chromatin accessibility and NKX2.2 binding

To determine the effect of *Klf4* misexpression on β cell gene expression we performed qPCR expression assays on several genes that also had associated chromatin changes in the *Klf4*-OE. Interestingly, there was an increase in *MafB* and *Irx2* expression in the *Klf4*-OE MIN6 cells (Supplemental Fig. S6A); however, there was no significant change in either *Ins2* or *Nkx6.1* expression, suggesting that KLF4 may be sufficient to activate α cell genes, but not to repress β cell genes in the context of β cells. Consistent with this, RNA-seq analysis of the *Klf4*-OE versus empty vector control MIN6 cells revealed that of the 2,005 genes dysregulated by *Klf4*-OE in β cells, 97% were upregulated genes. This included many important α cell enriched genes such as *Arx, MafB, Irx2, Gata6* and *Pyy* (Fig. 6D, Supplemental Data 6). Furthermore, comparison between *Klf4*-OE transcriptional dysregulation, cell-specific NKX2.2 binding, and *Klf4*-related chromatin changes revealed that in *Klf4*-OE β cells, 80 pro-α/anti-β loci were associated with significant changes in gene expression (Supplemental Data 7, Fig. 6E). Importantly, the majority of pro-α conditions of chromatin opening that were linked to changes in expression were located at proximal promoters and most anti-β conditions of chromatin closing that were linked to gene dysregulation were located at either introns or distal intergenic regions (Supplemental Fig. S6B; Fig. 6E).

Two striking examples of effects of *Klf4*-OE in β cells are at the proximal promoters of *MafB* and *Irx2*. In *Klf4*-OE β cells we observed a dramatic ATAC peak, signifying open chromatin accessibility, at the promoters *MafB* and *Irx2*, which was in sharp contrast to the absent or very low ATAC signal in the empty vector control (Fig. 6F,G). Furthermore, *MafB* and *Irx2* are normally expressed at low levels, if at all, in β cells, but in the *Klf4*-OE β cells we observed a significant increase in the expression of *MafB* and *Irx2* (Fig. 6F,G). These results demonstrate that *Klf4*-OE is sufficient to alter gene expression in β cells including the upregulation of α cell program genes and that many of these gene expression changes are linked to chromatin accessibility modulations that are conducive to an α cell specific NKX2.2 binding pattern.

### Klf4 misexpression in β cells is sufficient to induce changes in NKX2.2 binding

To assess whether misexpression of *Klf4* in MIN6 β cells stimulates changes in NKX2.2 binding, we assessed the DNA occupancy of KLF4-myc and NKX2.2 at the *MafB* and *Irx2* proximal promoters. This analysis revealed a significant increase in the mean enrichment of both KLF4-myc and NKX2.2 at the *MafB* promoter, but only a modest enrichment at the *Irx2* promoter in the *Klf4*-OE β cells compared to the empty vector control (Fig. 6H,J). However, linear regression analysis showed a significant positive correlation between KLF4 and NKX2.2 binding at the *MafB* and *Irx2* promoters, suggesting that an increase in KLF4 binding correlates to increased NKX2.2 binding (Supplemental Fig. S6C,D). These data together suggest that *Klf4* misexpression in MIN6 β cells is sufficient to increase NKX2.2 binding at the *MafB* and *Irx2* promoters–loci that are not normally bound by NKX2.2 in β cells–and induce the ectopic expression of these α cell genes.

## Discussion

In this study we sought to determine how the essential islet TF NKX2.2 exerted islet cell specific functions, with a focus on its role in regulating the α cell lineage. Deletion of *Nkx2*.2 in the α cell lineage resulted in a reduction of the α cell population size and dysregulation of islet hormone expression. Furthermore, as somewhat expected, transcriptomic analysis of *Nkx2.2^αΔ^* α cells identified that many transcriptional targets of NKX2.2 are α cell specific when compared to the β cell KO of NKX2.2. Surprisingly, however, we discovered that in α cells NKX2.2 binds more at proximal promoter regions compared to its binding in β cells and this cell-specific genome occupancy was at least partially facilitated by the reprogramming factor KLF4. This led to the characterization of KLF4 as an important islet α cell TF. Importantly, this study not only reveals the mechanism through which NKX2.2 can exert its islet cell specific activities, but it also identifies KLF4 as a novel islet α cell regulatory protein.

To assess the function of NKX2.2 in the α cell lineage, we created an *Nkx2.2* α cell KO mouse line using the highly specific and efficient *Gcg-iCre*. Although this is a useful model to study the role of NKX2.2 in maintaining α lineage identity, it does not address the α cell specific role of NKX2.2 in the specification of α cell identity or fate allocation.

However, to our knowledge there are no validated specific “pre-α” cell markers and associated Cre drivers that would allow us to currently address this question. Furthermore, due to the constitutive nature of the *Gcg-iCre*, we are unable to determine whether the observed phenotypes are due to a maturation and/or maintenance defect associated with the loss of NKX2.2 function. Future studies using an inducible Cre system to delete *Nkx2.2* from α cells after the islet maturation phase will discriminate between the role of NKX2.2 in maturation and maintenance.

Although there are striking molecular and morphometric phenotypes observed in the α cell KO of *Nkx2.2*, these defects only translate to a modest physiological phenotype. This is similar to many other mouse models that lack α cells or have dysfunctional α cells, including mice with deletions in *Arx, Pax6, MafB* and *Gcg* (Heller et al., 2004, Wilcox et al., 2013, Conrad et al., 2016). This discrepancy could be due to diminished necessity of α cells in mice compared to humans. Alternatively, there have been reports of the existence of extra-pancreatic sources of glucagon that could compensate for islet α cell dysfunctions (Sun et al., 2021).

Although these studies provide evidence that KLF4 is able to facilitate α cell specific binding of NKX2.2, not all of the NKX2.2 cell-specific targets can be accounted for by KLF4 cooperation. This suggests there may be additional cooperating factors that have not yet been characterized. We also identified motifs for the α-specific TF ETV1 that are enriched in NKX2.2-bound α cell promoters, supporting a role for ETV1 in facilitating NKX2.2 binding in α cells. Studies show that ETV1 regulates the transcriptome of many secretory and electrically-competent cell types, such as dopaminergic neurons and cardiomyocytes (Flames and Hobert, 2009, Shekhar et al., 2018). Although α cell enriched expression of ETV1 has been identified in several single cell RNA-seq studies, a functional role for ETV1 in the α cell population has yet to be described (Byrnes et al., 2018, Krentz et al., 2018).

Recent work in islet cells has revealed that many non-coding regulatory regions are simultaneously occupied by several islet TFs, demonstrating the extent of TF cooperation in islet gene regulation (Mawla et al., 2023). Interestingly, misexpression of *Klf4* in β cells both opened and closed chromatin. While it is easy to understand how KLF4 may open chromatin to facilitate NKX2.2 binding, given its role as a pioneer factor, it was surprising to observe a substantial number of closed chromatin regions at sites of β cell specific NKX2.2 binding. However, studies of KLF4 function in other systems have shown that KLF4 can open and close chromatin to regulate gene transcription (Moonen et al., 2022, Yu et al., 2021). This suggests that part of the role of KLF4 in promoting α cell identity is to create a chromatin landscape that deters β cell specific NKX2.2 occupancy. Furthermore, not all β cell specific NKX2.2 binding loci were affected by *Klf4* misexpression in β cells, which again suggests that other co-factors could be facilitating the β cell function of NKX2.2 and cooperating to promote β cell identity. Motif analysis of regions surrounding β cell NKX2.2 binding sites identified an enrichment of PDX1 and NKX6.1 motifs. PDX1 and NKX6.1—two essential β cell specific transcriptional regulators—are likely candidates for influencing NKX2.2 genome occupancy in β cells, much like KLF4 is directing the α cell specific promoter occupancy of NKX2.2.

In this study, we report for the first time that KLF4 promotes α cell identity by regulating cell-specific gene expression in cooperation with NKX2.2. Interestingly, several of the transcriptional targets of KLF4 do not overlap with NKX2.2 targets, suggesting that KLF4 plays a broader role maintaining α cell identity. α cells depleted of *Klf4* displayed many gene expression changes associated with known α cell specific genes. These data highlight the potency of KLF4 function alone as a pro-α cell factor that may be necessary and sufficient to promote α cell identity. However, additional studies will be needed to understand the breadth of *Klf4* function in α cell development, maturation and/or maintenance. Expression datasets of pancreatic tissue demonstrate that *Klf4* is highly enriched in the α cell population, however, *Klf4* RNA is undetectable during prenatal development suggesting that KLF4 likely functions after specification of the α lineage and potentially during maturation and/or maintenance of the α cells (Krentz et al., 2018).

In conclusion, we have demonstrated that NKX2.2 and KLF4 cooperate to maintain α cell identity in mice by regulating α cell specific gene expression. The functional cooperation between NKX2.2 and KLF4 emphasizes that the context in which a TF operates can be as important as its expression dynamics. Future studies into the role of TF cooperation in cell identity is especially important in light of the large-scale transcriptomic dysregulation that occurs during diabetic conditions (Brissova et al., 2018, Doliba et al., 2022, Dai et al., 2022, Avrahami et al., 2020). For example, loss of the appropriate complement of cell-type specific interacting factors could cause a TF to not only lose function but also gain function due to ectopically expressed interacting proteins and could explain the lineage transitions observed in islets from individuals with T1D or T2D. Our findings add significantly to the growing understanding of how TFs work together to regulate distinct islet identities and provide valuable insights that could be utilized in the study of diabetes etiology and stem cell therapies.

## Materials and Methods

### Generation of Nkx2.2 and Klf4 α cell knockout mice

Mice harboring the previously described *Nkx2.2^fl^* allele or the previously described *Klf4^fl^* allele were bred to *Gcg-iCre;Rosa26:tdTomato* mice (Mastracci et al., 2013b, ; Yoshida et al., 2008, Shiota et al., 2017, Madisen et al., 2010). PCR genotyping was performed to detect the *Nkx2.2^fl^, Gcg-iCre* and *tdTomato* alleles using the primers found in Supplemental Table 1.

#### Animal Maintenance

The mouse breeding colony was maintained on the C57B1/6J background. Mice were housed and treated in accordance with University of Colorado Institutional Animal Care and Use Committee approved protocols. Whenever possible both male and female mice were used in this study.

#### Immunofluorescence of fixed pancreas tissue sections

Pancreata were dissected, processed and stained as previously described (Gutierrez et al., 2017). Details provided in Supplemental Methods. Antibodies used in these experiments are listed in Supplemental Table 2. Imaging and morphometric image analyses are described in Supplemental methods.

#### Fluorescence Activated Cell Sorting (FACS) for murine α cells

Mouse Islets from mice with *Nkx2.2^fl/fl^;Gcg-iCre;Rosa26:tdTomato* and *Nkx2.2^+/+^;Gcg-iCre;Rosa26:tdTomato* genotypes were isolated from 5wk mice and dissociated into a single cell suspension via 0.25% trypsin/EDTA digestion. Trypsin digest was stopped by adding cold RPMI/10% FBS media to cell suspension. Cells were washed in cold 1x dPBS pH 7.1 twice before being strained through a cell-strainer tube cap. Cell suspensions were sorted on a Bio-Rad S3e cell sorter. Initial gating for all islets cells was done to avoid cell debris by excluding events with low forward scatter and low side scatter, adjusting the range of both settings to include the majority of events. Tomato+ cells were sorted directly into Qiagen RLT RNA lysis buffer. As an RNase inhibitor, β-mercaptoethanol was added to each tube of sorted cell lysate before being frozen down at -80°C for storage.

#### siRNA knockdown of gene expression in αTC α cells

Reverse transfection with siRNAs was performed on αTC cells to knockdown *Klf4* expression. 2.3 million αTC cells were seeded in a 60 mm cell culture treated plate and allowed to rest in the incubator for at least 30 minutes. Every plate was transfected with 35 μl of Invitrogen RNAiMax Lipofectamine (ThermoFisher 13778150) combined with either 350 pmol of Mouse *Klf4* ON-TARGETplus SMARTpool knockdown siRNA (Horizon L-040001-01-0010) or 350 pmol of siScramble ON-TARGETplus Non-targeting control pool (Horizon D-001810-10-05). Transfections were incubated for 48 h. Cell count and viability from these cell suspensions were done with Propidium Iodide on an Orflo Moxi GO II cell counter. RNA was extracted by the live cell samples to determine the efficiency of the knockdown via qPCR. Only samples with a 60% knockdown or greater were used in downstream experimentation.

#### Overexpression of myc-tagged constructs in islet cell culture

For *Klf4* overexpression (*Klf4*-OE) in the MIN6 and αTC cell lines, cells were transformed with p-GloMyc-Klf4 (AddGene 172870) and the empty vector control pcDNA-3.1 (Invitrogen V79020). 2.3×10^6^ cells were seeded in 60 mm cell culture plates in 4 mL media 24 h before transfection. Lipofectamine 3000 reagents (ThermoFisher L300008) were used to transfect 1 μg of plasmid. The cells were harvested 48 h post transfection. qPCR for *Klf4* was used to determine the level of *Klf4* expression over the empty vector control.

#### RNA-seq and differential gene expression analysis

Isolated total RNA samples were submitted to the Anschutz Genomics Core Facility. Only RNA with a RIN ≥ 8 were used for RNA-seq. Paired-end library preparation of RNA samples containing a total amount >500 ng was generated by the core using the Tecan Universal Plus mRNA-seq library preparation kit with NuQuant. Paired-end library preparation for samples containing <500 ng of RNA was generated by the core using the SMARTer Stranded Total RNA-seq kit for low input Ribo-depletion library preparation (Takara 634411). All RNA-seq libraries were run on an Illumina NovaSEQ 6000 or NovaSEQ X aiming for 40 million read pairs/80 million total reads per sample. Fastq output files were run through Fastqc (version 0.12.1) for quality control and adapters were trimmed from files using cutadapt (version 4.8) (Andrews, 2010, Martin, 2011). Alignment was performed using STAR (version 2.7.11b) to the to the mm10 genome before creating a counts table with featureCounts (version 2.0.6) (Dobin et al., 2013, Liao et al., 2014). DGE analysis was performed on RNA counts with experiments containing at least a sample size of 3 per condition, using DESeq2 or EdgeR programs in R (1.42.1, 4.0.16 and 4.3.2 version)(Love et al., 2013, Robinson et al., 2010). Significant gene expression change was determined with an adjusted p value or FDR cutoff of ≤ 0.05. RNA expression was visualized using the Integrated Genome Viewer (IGV).

#### ATAC-seq and differential chromatin accessibility analysis

Assay to detect transposase-accessible chromatin (ATAC) was performed to detect chromatin accessibility in *Klf4*-OE MIN6 cells. The Active Motif ATAC-seq Kit (Active Motif, 53150) was used to perform the ATAC and library prep with an initial live cell input of 100,000 according to manufacturer’s instructions. Libraries were sequenced on an Illumina NovaSEQ 6000 at a sequencing depth of 30 million read pairs/60 million total reads per sample. Output fastq files were sent through FastQC (version 0.12.1) for quality control and adapters were trimmed using cutadapt (version 4.8)(Andrews, 2010, Martin, 2011). Alignment to the mm10 genome was performed using bowtie2 (version 2.5.3) and chromatin peaks were called using MACS2 (version 2.2.9.1) peak caller with a q-value cutoff of ≤ 0.01 (Langmead and Salzberg, 2012, Feng et al., 2012). Differential chromatin accessibility analysis was performed with at least 3 biological replicates per condition using the Diffbind (version 3.12.0) software in R (version 4.3.2) at an FDR cutoff of ≤ 0.05. Chromatin peaks were annotated using ChIPseeker (version 1.38.0)(Ross-Innes et al., 2012, Wang et al., 2022). ATAC peaks were visualized using the IGV.

#### ChIP-seq and differential binding analysis

Chromatin immunoprecipitation (ChIP) was performed using the Cell Signaling Technologies SimpleChIP Enzymatic Chromatin IP Kit (Cell Signaling Technologies 9003S) according to manufacturer’s instructions. 10 μg of sheared chromatin per IP was incubated overnight at 4°C with 2 μg of anti-NKX2.2 antibody and 2 μg Normal Rabbit IgG antibody as a negative control. A 2% input sample was reserved and a positive control IP using the kit provided histone HS antibody was initially performed to determine kit efficacy. Immuno-enriched ChIP samples were purified via magnetic beads and their concentration was determine via Nanodrop. qPCR was used to determine enrichment of experimental ChIP samples over IgG at positive control loci before sending samples to the Anschutz Genomics Core for quality and sequencing. ChIP libraries were prepared by the University of Colorado Anschutz Genomics core using the Tecan Ovation Ultralow System V2 and then sequenced on a NovaSEQ 6000. ChIP-seq read depth was 10 million read pairs/20 million total reads. Output fastq files were run through fastQC (version 0.12.1) for quality control and adapters were trimmed using cutadapt (version 4.8(Andrews, 2010, Martin, 2011). Alignment to the mm10 genome was performed using bowtie2 and binding peaks were called compared to the input samples by using MACS2 peak caller (version 2.2.9.1) with a q-value cutoff of ≤ 0.01. ChIP peaks were annotated using the ChIPseeker software (version 1.38.0) (Langmead and Salzberg, 2012, Feng et al., 2012, Yu et al., 2015). When appropriate, differential binding analysis was performed using the csaw program (version 1.36.1) in R (version 4.3.2) on 3 or more replicates per condition. ChIP peaks were visualized using IGV.(Lun and Smyth, 2016).

#### Cut&Run binding assay

Cleavage Under Targets and Release Using Nuclease (Cut&Run) assays to detect DNA binding of NKX2.2 and KLF4 were conducted using the Cell Signaling Technologies Cut&Run Assay Kit (Cell Signaling Technology, 86652S). 100,000 live cells per Cut&Run preparation were fixed using 0.1% formaldehyde for 2 minutes. Fixed cells were incubated for 2 h at 4°C with Concanavalin A beads, and the anti-NKX2.2 antibody, anti-KLF4 antibody, IgG XP Isotype control antibody or H3K4me3 positive control antibody. Cell/bead suspensions were then incubated with the pAG-MNase enzyme for 1 h at 4°C before activation of the MNase with CaCl2 and incubation at 4°C for 30 minutes. A 5 ng Yeast Sample Normalization Spike-In was added to each Cut&Run reaction. 100,000 fixed cells were also used to generate the input sample.

#### Quantification of binding assay enrichment via qPCR

Locus specific enrichment in ChIP and Cut&Run samples due to TF occupancy was measured via qPCR. Primers were designed to amplify 60-80bp regions, to have an optimal melting temperature of 60c and have a GC content close to 50%. Cell signaling Technologies SimpleChIP Universal qPCR Master Mix (Cell Signaling Technologies 88989P) was used to amplify genomic loci on a Bio-Rad CFX96 Real-Time Machine using the standard SimpleChIP PCR program. For Cut&Run qPCR, reactions that included primers for amplifying the spike-in normalization locus were included in every experiment. Cut&Run sample enrichment for each assayed locus was calculated via the percent input method and normalized to the lowest spike-in value. Percent input values of experimental Cut&Run samples were compared to the percent input of the IgG negative control. ChIP sample enrichment for each assayed locus was calculated using the percent input method but with no spike-in normalization. Enrichment at a ChIP sample locus was determined by comparing the percent input of the experimental sample to the percent input of the IgG.

#### Gene set enrichment analysis

We calculated all gene set enrichments using a hypergeometric test, e.g. if the frequency of genes related to a defined group are statistically higher in our datasets than background frequency. Enrichment of α, β or δ cell specific genes in our datasets were calculated manually by defining our own islet expression gene sets. We determined the number of cell-enriched genes present in the islet transcriptome using the Huising Lab’s published islet RNA-seq dataset and pairwise EdgeR (version 4.016) DGE analyses (α vs β, α vs δ, or β vs δ). We determined α, β, or δ cell-enriched genes as being ≤ -1 log2 foldchange or ζ 1 log2 foldchange. This list of cell-enriched islet genes was then used to annotate the genes present in our datasets as cell-enriched. The proportion of cell-enriched genes in our dataset was statistically compared to the proportion of cell-enriched genes in the entire islet transcriptome using a hypergeometric test. Our p-value cutoff for islet cell expression enrichment was ≤ 0.01.

#### TF motif enrichment analysis

The enrichment of TF motifs was performed using the α cell NKX2.2 ChIP-seq generated in this study and the β cell NKX2.2 ChIP-seq generated in a previously published study. We performed a non-biased search for motif enrichment using the HOMER2 motif enrichment software (version v5) and the homer motif database, where the α and β cell NKX2.2 ChIP-seqs were analyzed separately (Heinz et al., 2010). Islet cell specific expression of TFs associated with enriched motifs was determined using the Huising Lab’s previously published islet RNA-seq DGE analysis described above. The islet cell expression annotation was used to determine which of the enriched motifs are specifically expressed in α or β cells. The top α cell specific motif candidates were then used for motif enrichment analysis using TFBSTools, universalmotif and seqLogo R software (versions 1.40.0, 1.20.0, 1.68.0)(Tan and Lenhard, 2016, Tremblay, 2021, Bembom, 2007). Motifs were retrieved from either the homer motif database or the JASPAR database (https://jaspar.elixir.no/, http://homer.ucsd.edu/homer/). Motifs were searched one at a time for presence within +/-100bp of the α or β-specific NKX2.2 binding summits with a minimum matching score of 80%. Enrichment of each candidate motif within each genomic feature subset (promoter <= 1 kb, other intron, distal intergenic, etc) was determined by comparing the proportion of motifs found in the NKX2.2 binding site subsets to the proportion of motifs found in 10,000 random genomic sequences using a hypergeometric test. Enrichment p-value cutoff was set at ≤ 0.01 for motif analyses.

### Statistical analysis

T-tests, Z-tests, ANOVA with Tukey’s post hoc, and hypergeometric tests were performed using the R programing language (version 4.3.2). Cutoffs for p-values, FDR, adjusted-p and q-values were set no higher than ≤ 0.05. Multiple comparisons adjustments were performed only if the number of tests was greater than 10 in a single analysis. When appropriate, a minimum of 3 biological replicates were used in statistical analysis.

## Data Availability

All sequencing data generated in this study have been uploaded to the Gene Expression Omnibus (GEO) under the accession numbers: GSE273739, GSE273740, GSE273741, GSE273742, GSE273743, GSE273744, and GSE273745.

## Acknowledgements

We thank all past and current members of the Sussel laboratory for helpful conversations and reading of this manuscript. We thank K. Scott Beard in the Barbara Davis Center Islet Core for islet isolations used in this study. We are grateful to the University of Colorado Anschutz Medical Campus Cancer Center Genomics Shared Resource Core Facility (RRID:SCR_021984) for library preparation and next generation sequencing. We are thankful for critical review of this manuscript that was in part done by students of the Cells, Stem cells and Developmental Biology Graduate Program’s Advanced Writing Workshop. Funding for this work was provided by the National Institute for Health grants (F31 DK13028 to E.P.B.; R01 DK082590, R01 DK118155, and U01 DK127505 to LS. Core facility support was provided by the University of Colorado Diabetes Center P30 DK116073.)

## Author contributions

E.P.B. and L.S. conceived the project. E.P.B. and M.R.C. performed the majority of the experiments and T.L. and M.V.O. contributed with minor experiments. Bioinformatics analysis was conducted by E.P.B. and K.L.W. E.P.B. and L.S. cowrote the paper including input from all authors.

## References

1. Andrews, S. 2010. FastQC: a quality control tool for high throughput sequence data. [Online]. Available: http://www.bioinformatics.babraham.ac.uk/projects/fastqc [Accessed].

2. Arnes, L., Leclerc, K., Friel, J. M., Hipkens, S. B., Magnuson, M. A. & Sussel, L. 2012. Generation of Nkx2. 2: lacZ mice using recombination-mediated cassette exchange technology. Genesis, 50, 612–624.

3. Artner, I. L. J.; Hang, Y.; Elghazi, L.; Chisler, J.; Henderson, E.; Sosa-Pineda, B.; Stein, R. 2006. MAFB: An Activator of the Glucagon Gene Expressed in Developing Islet Alpha- and Beta-Cells. Diabetes, 55.

4. Avrahami, D., Wang, Y. J., Schug, J., Feleke, E., Gao, L., Liu, C., Consortium, H., Naji, A., et al. 2020. Single-cell transcriptomics of human islet ontogeny defines the molecular basis of beta-cell dedifferentiation in T2D. Mol Metab, 42, 101057.

5. Bembom, O. 2007. Sequence logos for DNA sequence alignments. R package version, 1.

6. Bishop, C. D., Macneil, K. E., Patel, D., Taylor, V. J. & Burke, R. D. 2013. Neural development in Eucidaris tribuloides and the evolutionary history of the echinoid larval nervous system. Developmental biology, 377, 236–244.

7. Bolli, G., Feo, P. D., Compagnucci, P., Cartechini, M. G., Angeletti, G., Santeusanio, F., Brunetti, P. & Gerich, J. E. 1983. Abnormal Glucose Counterregulation in Insulin-dependent Diabetes Mellitus: Interaction of AnI-Insulin Antibodies and Impaired Glucagon and Epinephrine Secretion. Diabetes, 32, 134–141.

8. Brissova, M., Haliyur, R., Saunders, D., Shrestha, S., Dai, C., Blodgett, D. M., Bottino, R., Campbell-Thompson, M., et al. 2018. Alpha Cell Function and Gene Expression Are Compromised in Type 1 Diabetes. Cell Rep, 22, 2667–2676.

9. Brooks, E. P. & Sussel, L. 2023. Not the second _ddle: ± cell development, identity, and function in health and diabetes. Journal of Endocrinology, 258.

10. Byrnes, L. E., Wong, D. M., Subramaniam, M., Meyer, N. P., Gilchrist, C. L., Knox, S. M., Tward, A. D., Ye, C. J., et al. 2018. Lineage dynamics of murine pancreatic development at single-cell resolution. Nat Commun, 9, 3922.

11. Chang, Y.-H., Katoh, M. C., Abdellatif, A. M., Xiafukaiti, G., Elzeftawy, A., Ojima, M., Mizuno, S., Kuno, A., et al. 2020. Uncovering the role of MAFB in glucagon production and secretion in pancreatic ±-cells using a new ±-cell-specific Mal conditional knockout mouse model. Experimental Animals, 69, 178–188.

12. Chung, C. H., Hao, E., Piran, R., Keinan, E. & Levine, F. 2010. Pancreatic beta-cell neogenesis by direct conversion from mature alpha-cells. Stem Cells, 28, 1630–8.

13. Churchill, A. J., Gutierrez, G. D., Singer, R. A., Lorberbaum, D. S., Fischer, K. A. & Sussel, L. 2017. Genetic evidence that Nkx2.2 acts primarily downstream of Neurog3 in pancreatic endocrine lineage development. Elife, 6.

14. Collombat, P., Hecksher-Sorensen, J., Krull, J., Berger, J., Riedel, D., Herrera, P. L., Serup, P. & Mansouri, A. 2007. Embryonic endocrine pancreas and mature beta cells acquire alpha and PP cell phenotypes upon Arx misexpression. J Clin Invest, 117, 961–70.

15. Conrad, E., Dai, C., Spaeth, J., Guo, M., Cyphert, H. A., Scoville, D., Carroll, J., Yu, W. M., et al. 2016. The MAFB transcription factor impacts islet alpha-cell function in rodents and represents a unique signature of primate islet beta-cells. Am J Physiol Endocrinol Metab, 310, E91–E102.

16. Courtney, M., Gjernes, E., Druelle, N., Ravaud, C., Vieira, A., Ben-Othman, N., Pfeifer, A., Avolio, F., et al. 2013. The inactivation of Arx in pancreatic alpha-cells triggers their neogenesis and conversion into functional beta-like cells. PLoS Genet, 9, e1003934.

17. Dai, X. Q., Camunas-Soler, J., Briant, L. J. B., Dos Santos, T., Spigelman, A. F., Walker, E. M., Arrojo, E. D. R., Bautista, A., et al. 2022. Heterogenous impairment of alpha cell function in type 2 diabetes is linked to cell maturation state. Cell Metab, 34, 256–268 e5.

18. Di Giammartino, D. C., Kloetgen, A., Polyzos, A., Liu, Y., Kim, D., Murphy, D., Abuhashem, A., Cavaliere, P., et al. 2019. KLF4 is involved in the organization and regulation of pluripotency-associated three-dimensional enhancer networks. Nature cell biology, 21, 1179–1190.

19. Digruccio, M. R., Mawla, A. M., Donaldson, C. J., Noguchi, G. M., Vaughan, J., Cowing-Zitron, C., Van Der Meulen, T. & Huising, M. O. 2016. Comprehensive alpha, beta and delta cell transcriptomes reveal that ghrelin selectively activates delta cells and promotes somatostatin release from pancreatic islets. Mol Metab, 5, 449–458.

20. Dobin, A., Davis, C. A., Schlesinger, F., Drenkow, J., Zaleski, C., Jha, S., Batut, P., Chaisson, M., et al. 2013. STAR: ultrafast universal RNA-seq aligner. BioinformaLcs, 29, 15–21.

21. Doliba, N. M., Rozo, A. V., Roman, J., Qin, W., Traum, D., Gao, L., Liu, J., Manduchi, E., et al. 2022. alpha Cell dysfunction in islets from nondiabetic, glutamic acid decarboxylase autoantibody-positive individuals. J Clin Invest, 132.

22. Doyle, M. J., Loomis, Z. L. & Sussel, L. 2007. Nkx2.2-repressor activity is suffcient to specify alpha-cells and a small number of beta-cells in the pancreatic islet. Development, 134, 515–23.

23. Doyle, M. J. & Sussel, L. 2007. Nkx2.2 regulates beta-cell function in the mature islet. Diabetes, 56, 1999–2007.

24. Feng, J., Liu, T., Qin, B., Zhang, Y. & Liu, X. S. 2012. Identifying ChIP-seq enrichment using MACS. Nature protocols, 7, 1728–1740.

25. Flames, N. & Hobert, O. 2009. Gene regulatory logic of dopamine neuron differentiation. Nature, 458, 885–889.

26. Gerich, J. E., Langlois, M., Noacco, C., Karam, J. H. & Forsham, P. H. 1973. Lack Of Glucagon Response To Hypoglycemia In Diabetes: Evidence For An Intrinsic Pancreaic Alpha Cell Defect. Science, 182, 171–173.

27. Gosmain, Y., Avril, I., Mamin, A. & Philippe, J. 2007. Pax-6 and c-Maf functionally interact with the alpha-cell-specific DNA element G1 in vivo to promote glucagon gene expression. J Biol Chem, 282, 35024–34.

28. Gosmain, Y., Marthinet, E., Cheyssac, C., Guerardel, A., Mamin, A., Katz, L. S., Bouzakri, K. & Philippe, J. 2010. Pax6 controls the expression of critical genes involved in pancreatic {alpha} cell differentiation and function. J Biol Chem, 285, 33381–33393.

29. Gutierrez, G. D., Bender, A. S., Cirulli, V., Mastracci, T. L., Kelly, S. M., Tsirigos, A., Kaestner, K. H. & Sussel, L. 2017. Pancreatic beta cell identity requires continual repression of non-beta cell programs. J Clin Invest, 127, 244–259.

30. Heinz, S., Benner, C., Spann, N., Bertolino, E., Lin, Y. C., Laslo, P., Cheng, J. X., Murre, C., et al. 2010. Simple combinations of lineage-determining transcription factors prime cis-regulatory elements required for macrophage and B cell idenIIes. Molecular cell, 38, 576–589.

31. Heller, R. S., Stoffers, D. A., Liu, A., Schedl, A., Crenshaw Iii, E. B., Madsen, O. D. & Serup, P. 2004. The role of Brn4/Pou3f4 and Pax6 in forming the pancreatic glucagon cell identity. Developmental biology, 268, 123–134.

32. Katoh, M. C., Jung, Y., Ugboma, C. M., Shimbo, M., Kuno, A., Basha, W. A., Kudo, T., Oishi, H., et al. 2018. MafB Is Critical for Glucagon Production and Secretion in Mouse Pancreatic alpha Cells In Vivo. Mol Cell Biol, 38.

33. Krentz, N. A., Lee, M. Y., Xu, E. E., Sproul, S. L., Maslova, A., Sasaki, S. & Lynn, F. C. 2018. Single-cell transcriptome profiling of mouse and hESC-derived pancreatic progenitors. Stem cell reports, 11, 1551–1564.

34. Langmead, B. & Salzberg, S. L. 2012. Fast gapped-read alignment with Bowtie 2. Nature methods, 9, 357–359.

35. Liao, Y., Smyth, G. K. & Shi, W. 2014. featureCounts: an efficient general purpose program for assigning sequence reads to genomic features. BioinformaLcs, 30, 923–930.

36. Love, M., Anders, S. & Huber, W. 2013. Differential analysis of RNA-Seq data at the gene level using the DESeq2 package. Heidelberg: European Molecular Biology Laboratory (EMBL).

37. Lun, A. T. & Smyth, G. K. 2016. csaw: a Bioconductor package for differential binding analysis of ChIP-seq data using sliding windows. Nucleic acids research, 44, e45.

38. Maclean, N. & Ogilvie, R. F. 1959. Observaions On The Pancreaic Islet Issue Of Young Diabeic Subjects. Diabetes, 8, 83–91.

39. Madisen, L., Zwingman, T. A., Sunkin, S. M., Oh, S. W., Zariwala, H. A., Gu, H., Ng, L. L., Palmiter, R. D., et al. 2010. A robust and high-throughput Cre reporting and characterization system for the whole mouse brain. Nature neuroscience, 13, 133–140.

40. Marchetti, P., Dotta, F., Ling, Z., Lupi, R., Del Guerra, S., Santangelo, C., Realacci, M., Marselli, L., et al. 2000. Function of pancreatic islets isolated from a type 1 diabetic patient. Diabetes Care, 23, 701–3.

41. Martin, M. 2011. Cutadapt removes adapter sequences from high-throughput sequencing reads. EMBnet. journal, 17, 10–12.

42. Mastracci, T. L., Anderson, K. R., Papizan, J. B. & Sussel, L. 2013a. Regulation of Neurod1 contributes to the lineage potential of Neurogenin3+ endocrine precursor cells in the pancreas. PLoS Genet, 9, e1003278.

43. Mastracci, T. L., Lin, C. S. & Sussel, L. 2013b. Generation of mice encoding a conditional allele of Nkx2.2. Transgenic Res, 22, 965–72.

44. Mastracci, T. L., Wilcox, C. L., Arnes, L., Panea, C., Golden, J. A., May, C. L. & Sussel, L. 2011. Nkx2.2 and Arx genetically interact to regulate pancreatic endocrine cell development and endocrine hormone expression. Dev Biol, 359, 1–11.

45. Mawla, A. M., Van Der Meulen, T. & Huising, M. O. 2023. Chromatin accessibility diferences between alpha, beta, and delta cells identifies common and cell type-specific enhancers. BMC genomics, 24, 1–22.

46. Moonen, J.-R., Chappell, J., Shi, M., Shinohara, T., Li, D., Mumbach, M. R., Zhang, F., Nair, R. V., et al. 2022. KLF4 recruits SWI/SNF to increase chromatin accessibility and reprogram the endothelial enhancer landscape under laminar shear stress. Nature communications, 13, 4941.

47. Mourao, L., Zeeman, A. L., Wiese, K. E., Bongaarts, A., Oudejans, L. L., Martinez, I. M., Van De Grift, Y. B., Jonkers, J., et al. 2021. Hyperactive WNT/CTNNB1 signaling induces a competing cell proliferation and epidermal differentiation response in the mouse mammary epithelium. bioRxiv, 2021.06.22.449461.

48. Müller, W. A., Faloona, G. R., Aguilar-Parada, E. & Unger, R. H. 1970. Abnormal Alpha-Cell Funcion In Diabetes. New England Journal Of Medicine, 283, 109–115.

49. Nakagawa, M., Koyanagi, M., Tanabe, K., Takahashi, K., Ichisaka, T., Aoi, T., Okita, K., Mochiduki, Y., et al. 2008. Generation of induced pluripotent stem cells without Myc from mouse and human fibroblasts. Nature biotechnology, 26, 101–106.

50. Nishimura, W., Kondo, T., Salameh, T., El Khattabi, I., Dodge, R., Bonner-Weir, S. & Sharma, A. 2006. A switch from MafB to MafA expression accompanies differentiation to pancreatic β-cells. Developmental biology, 293, 526–539.

51. Papizan, J. B., Singer, R. A., Tschen, S. I., Dhawan, S., Friel, J. M., Hipkens, S. B., Magnuson, M. A., Bhushan, A., et al. 2011. Nkx2.2 repressor complex regulates islet beta-cell specification and prevents beta-to-alpha-cell reprogramming. Genes Dev, 25, 2291–305.

52. Pictet, R. L., Clark, W. R., Williams, R. H. & Rutter, W. J. 1972. An ultrastructural analysis of the developing embryonic pancreas. Developmental biology, 29, 436–467.

53. Prasadan, K., Tulachan, S., Guo, P., Shiota, C., Shah, S. & Gittes, G. 2010. Endocrine-committed progenitor cells retain their differentiation potential in the absence of neurogenin-3 expression. Biochemical and biophysical research communicaLons, 396, 1036–1041.

54. Robinson, M. D., Mccarthy, D. J. & Smyth, G. K. 2010. edgeR: a Bioconductor package for differential expression analysis of digital gene expression data. bioinformaLcs, 26, 139–140.

55. Rosenzweig, J. M., Glenn, J. D., Calabresi, P. A. & Whartenby, K. A. 2013. KLF4 modulates expression of IL-6 in dendritic cells via both promoter activation and epigenetic modification. Journal of Biological Chemistry, 288, 23868–23874.

56. Ross-Innes, C. S., Stark, R., Teschendorff, A. E., Holmes, K. A., Ali, H. R., Dunning, M. J., Brown, G. D., Gojis, O., et al. 2012. Differential oestrogen receptor binding is associated with clinical outcome in breast cancer. Nature, 481, 389–393.

57. Sander, M., Neubüser, A., Kalamaras, J., Ee, H. C., Martin, G. R. & German, M. S. 1997. Geneic Analysis Reveals That Pax6 Is Required For Normal Transcripion Of Pancreaic Hormone Genes And Islet Development. Genes & Development, 11, 1662–1673.

58. Schaum, N., Karkanias, J., Neff, N. F., May, A. P., Quake, S. R., Wyss-Coray, T., Darmanis, S., Batson, J., et al. 2018. Single-cell transcriptomics of 20 mouse organs creates a Tabula Muris. Nature, 562, 367–372.

59. Schindelin, J., Arganda-Carreras, I., Frise, E., Kaynig, V., Longair, M., Pietzsch, T., Preibisch, S., Rueden, C., et al. 2012. Fiji: an open-source plauorm for biological-image analysis. Nature methods, 9, 676-682.

60. Shekhar, A., Lin, X., Lin, B., Liu, F.-Y., Zhang, J., Khodadadi-Jamayran, A., Tsirigos, A., Bu, L., et al. 2018. ETV1 activates a rapid conduction transcriptional program in rodent and human cardiomyocytes. ScienL_c reports, 8, 9944.

61. Shiota, C., Prasadan, K., Guo, P., Fusco, J., Xiao, X. & Gittes, G. K. 2017. Gcg (CreERT2) knockin mice as a tool for genetic manipulation in pancreatic alpha cells. Diabetologia, 60, 2399–2408.

62. Singer, R. A., Arnes, L., Cui, Y., Wang, J., Gao, Y., Guney, M. A., Burnum-Johnson, K. E., Rabadan, R., et al. 2019. The Long Noncoding RNA Paupar Modulates PAX6 Regulatory Activities to Promote Alpha Cell Development and Function. Cell Metab, 30, 1091–1106 e8.

63. Smith, S. B., Watada, H. & German, M. S. 2004. Neurogenin3 activates the islet differentiation program while repressing its own expression. Molecular Endocrinology, 18, 142–149.

64. Soufi, A., Donahue, G. & Zaret, K. S. 2012. Facilitators and impediments of the pluripotency reprogramming factors’ initial engagement with the genome. Cell, 151, 994–1004.

65. Sun, E. W., Martin, A. M., De Fontgalland, D., Sposato, L., Rabbitt, P., Hollington, P., Wattchow, D. A., Colella, A. D., et al. 2021. Evidence for glucagon secretion and function within the human gut. Endocrinology, 162, bqab022.

66. Sussel, L. K. J.; Hartigan-O’Connor, D. J.; Meneses, J. J.; Pedersen, R. A.; Rubenstein, J. L. R. And German, M. S. 1998. Mice Lacking The Homeodomain Transcripion Factor Nkx2.2 have diabetes due to arrested differentiation of pancreatic ² cells. Development, 125, 2213–2221.

67. Swisa, A., Avrahami, D., Eden, N., Zhang, J., Feleke, E., Dahan, T., Cohen-Tayar, Y., Stolovich-Rain, M., et al. 2017. PAX6 maintains beta cell identity by repressing genes of alternative islet cell types. J Clin Invest, 127, 230–243.

68. Takahashi, K. & Yamanaka, S. 2006. Induction of pluripotent stem cells from mouse embryonic and adult _broblast cultures by defined factors. cell, 126, 663–676.

69. Tan, G. & Lenhard, B. 2016. TFBSTools: an R/bioconductor package for transcription factor binding site analysis. BioinformaLcs, 32, 1555–1556.

70. Tremblay, B. 2021. universalmotif: import, modify, and export motifs with R. R package version 1.8. 2.

71. Wang, Q., Li, M., Wu, T., Zhan, L., Li, L., Chen, M., Xie, W., Xie, Z., et al. 2022. Exploring epigenomic datasets by ChIPseeker. Current Protocols, 2, e585.

72. Wei, Z., Gao, F., Kim, S., Yang, H., Lyu, J., An, W., Wang, K. & Lu, W. 2013. Klf4 organizes long-range chromosomal interactions with the oct4 locus in reprogramming and pluripotency. Cell stem cell, 13, 36–47.

73. Wilcox, C. L., Terry, N. A., Walp, E. R., Lee, R. A. & May, C. L. 2013. Pancreatic alpha-cell specific deletion of mouse Arx leads to alpha-cell identity loss. PLoS One, 8, e66214.

74. Yoshida, T., Kaestner, K. H. & Owens, G. K. 2008. Conditional deletion of Kruppel-like factor 4 delays downregulation of smooth muscle cell differentiation markers but accelerates neointimal formation following vascular injury. CirculaLon research, 102, 1548–1557.

75. Yu, G., Wang, L.-G. & He, Q.-Y. 2015. ChIPseeker: an R/Bioconductor package for ChIP peak annotation, comparison and visualization. BioinformaLcs, 31, 2382–2383.

76. Yu, S., Guo, J., Sun, Z., Lin, C., Tao, H., Zhang, Q., Cui, Y., Zuo, H., et al. 2021. BMP2-dependent gene regulatory network analysis reveals Klf4 as a novel transcription factor of osteoblast differentiation. Cell death & disease, 12, 197.

